# RpoS Contributes to Successful Opportunistic Colonization by Human Enteric Pathogens during Plant Disease

**DOI:** 10.1101/2020.11.24.397208

**Authors:** Amelia H. Lovelace, Sangwook Lee, Diana M. Downs, Ziad Soufi, Pedro Bota, Gail M. Preston, Brian H. Kvitko

**Author notes:** Address correspondence to Brian H. Kvitko,.

## Abstract

With an increase in foodborne illnesses associated with the consumption of fresh produce, it is important to understand the interactions between human bacterial enteric pathogens and plants. It was previously established that diseased plants can create a permissive environment for opportunistic endophytic colonization of enteric pathogens. However, the factors that contribute to the colonization of enteric pathogens during plant disease are largely unknown. Here, we show that both strain and plant host factors contribute to significantly increased populations of enteric pathogens when co-inoculated with the plant pathogen, *P. syringae* pv. *tomato*. The two *Salmonella enterica* strains DM10000 and 14028S, differ in their ability to metabolize host-derived apoplastic carbohydrates dependent on the sigma factor RpoS. The *rpoS* gene is an important strain factor for endophytic colonization by *S. enterica* during plant disease. Our results suggest that *rpoS* plays a crucial role during *in planta* colonization, balancing nutrient metabolism and stress responses.

**Importance:** Foodborne illnesses caused by the bacterial human enteric pathogens, *E. coli* O157:H7 and *S. enterica*, often results in vomiting and diarrhea. If left untreated, this illness can cause dehydration and sometimes death of a patient. Both *E. coli* O157:H7 and *S. enterica* have caused repeated fresh produce-associated epidemics. Crop disease could promote the ability of plants to act as reservoirs for produce-borne outbreaks. Plant pathogens dampen plant immunity, which allows for a more permissive environment for human enteric pathogens to grow. These internalized enteric pathogen populations are especially dangerous since they cannot be removed by washing alone. Therefore, the need to understand the factors that contribute to the opportunistic colonization of human enteric pathogens during plant disease is apparent. Our research has identified host and strain factors that contribute to opportunistic colonization of diseased plants, which will inform the development of future management strategies to mitigate future outbreaks.

## Introduction

Fresh produce has become an increasing source for foodborne illnesses. In the United States, fresh produce foodborne outbreaks have increased from 0.7% of reported outbreaks in the 1970s to 33% in 2011 (1). The cause of this increase is largely unknown. The two most prevalent bacterial human pathogens causing fresh-produce-associated epidemics of enteric illnesses include *Salmonella enterica,* which caused 48% of such outbreaks from 1973 to 1997, and Shiga-toxin producing *Escherichia coli* (2). In 2019, there were two *E. coli* O157:H7 outbreaks on romaine lettuce representing the largest *E. coli* flare-up in more than a decade and causing more than 200 illnesses and five deaths (3). Fresh produce can therefore serve as a vector for these bacterial enteric pathogens to enter human hosts (4, 5).

Research efforts to understand produce-borne illness have typically focused on the attachment, fitness, and persistence of epiphytic colonization of human enteric pathogens on fresh produce (6, 7). Endophytic colonization of *S. enterica* serovar Typhimurium in tomato and *E. coli* O157:H7 in lettuce makes these pathogens challenging to remove by surface-sanitization treatments (8, 9). Thus endophytic populations of human pathogens on produce could pose a significant public health concern. The underlying genetic factors that contribute to the endophytic colonization of plants by human enteric pathogens are generally not well understood.

Under natural conditions, the enteric populations within plant tissue may be restricted from reaching a human infectious dose due to the robust innate immunity of plants (10). Plants possess diverse surface receptor proteins termed Pattern Recognition Receptors (PRRs) that detect non-adapted microbes by binding to conserved “non-self” microbe-associated molecular patterns (MAMPs) such as bacterial flagellin, lipopolysaccharides, and peptidoglycan (11–13). The MAMPs found in both *Salmonella* and Shiga toxin-producing *E. coli* can be detected by plant PRRs (14, 15). This recognition event induces an immune response called Pattern Triggered Immunity (PTI) that restricts colonization by diverse microbes (non-adapted microbes) (16). Well-adapted plant pathogenic bacterial species, such as *Pseudomonas syringae*, have evolved active mechanisms to suppress PTI (17). To do so, hemibiotrophic bacterial plant pathogens deliver virulence-associated proteins termed effectors into the plant cell through the Type III Secretion System (T3SS). These effectors modify plant host cell targets and suppress PTI-associated immune signaling (18). Effectors play multiple roles in creating a permissive environment for bacterial proliferation by also manipulating host systems to release water and presumably nutrients into the apoplast (19, 20). Unlike hemibiotrophic organisms that require living plant cells to complete their disease cycle, necrotrophic bacterial plant pathogens such as those that cause soft rot, enzymatically degrade and metabolize plant cell wall polysaccharides once populations reach a quorum (21). This creates a permissive environment for bacterial proliferation by releasing cellular contents that can serve as substrates for growth.

The permissive environment produced by plant pathogenic bacteria through the delivery of immune dampening effectors or cell wall degrading enzymes can affect the other bacterial organisms associated with these plant pathogens It has been demonstrated that *S. enterica* can benefit from the environments established by plant pathogens of various lifestyles and can grow to higher populations in the presence of a compatible plant pathogen (22–25). Similarly, *E. coli* O157:H7 increases in population size when co-inoculated with a soft rot bacterial plant pathogen on lettuce (26). Consequently, diseased plants can potentially serve as important reservoirs for human enteric pathogens colonizing the internal tissues of fresh produce. However, the factors important for endophytic colonization of plants by human enteric pathogens under diseased conditions established by plant pathogens are largely unknown.

Here, we present the results from a co-inoculation analysis of the human enteric pathogens, *S. enterica* and *E. coli* O157:H7, with and without the plant pathogen *P. syringae* pv. *tomato* DC3000 in three compatible plant hosts, *Arabidopsis thaliana*, *Nicotiana benthamiana*, and collard (*Brassica oleracea* var. *acephala*). Two genotypes of *P. syringae* were used in the co-inoculation study, one with a functional T3SS and one without a functional T3SS to generate a permissive and non-permissive environment for bacterial colonization respectively. Our study reveals that human enteric pathogens do not benefit from the permissive environment established by *P. syringae* in all cases and both strain and host factors contribute to their opportunistic colonization of host leaf tissue during plant disease. Based on natural variation between *S. enterica* strains, RpoS was identified as a factor important for *S. enterica* to metabolize carbohydrates present within the plant apoplast. This sigma factor also plays a role in *in planta* colonization by enteric pathogens during plant disease, most likely by modulating bacterial stress responses.

## Results

### Increased endophytic colonization of *E. coli* O157:H7 during plant disease established by *P. syringae* is dependent on both *E. coli* and *P. syringae* initial populations

The bacterial pathogen *P. syringae* pv. *tomato* DC3000 (DC3K) has a well-studied repertoire of virulence factors allowing it to infect multiple model plant hosts, including *A. thaliana* and *N. benthamiana* (In the absence of recognition of HopQ1-1 effector by the immune receptor Roq1). In addition, the PTI response is well-characterized in these two hosts making these model plants perfect for elucidating the factors important for endophytic human enteric pathogen colonization during DC3K infection. Co-inoculation assays were performed on adult *A. thaliana* and *N. benthamiana* plants by syringe infiltration using two compatible DC3K strains (DC3K and DC3KΔ*hopQ1-1*) with a functional T3SS (T3SS+) or a DC3K strain without a functional T3SS (DC3KΔ*hrcC*, T3SS-) and the human enteric pathogen, non-toxigenic *E. coli* O157:H7 5-11 (O157:H7) (Table 1). To determine the ratio of O157:H7 to DC3K required for permissive or non-permissive growth in these two hosts, a range of starting inoculum concentrations were tested.

**Table 1.**
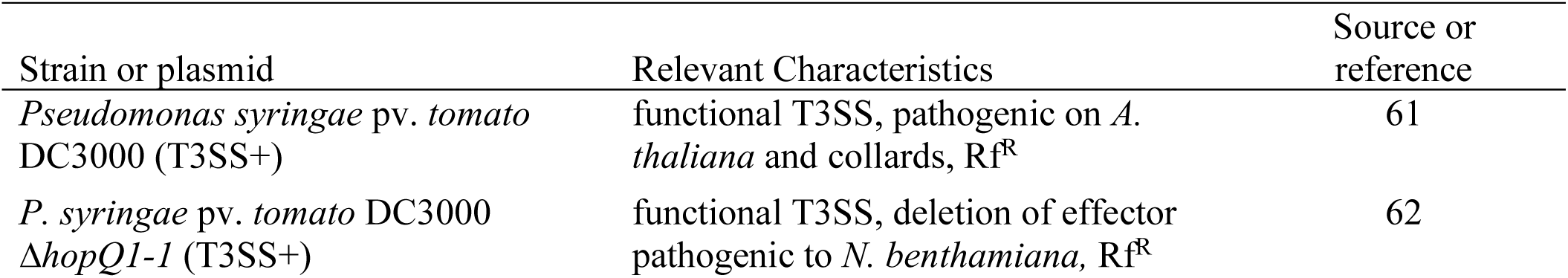
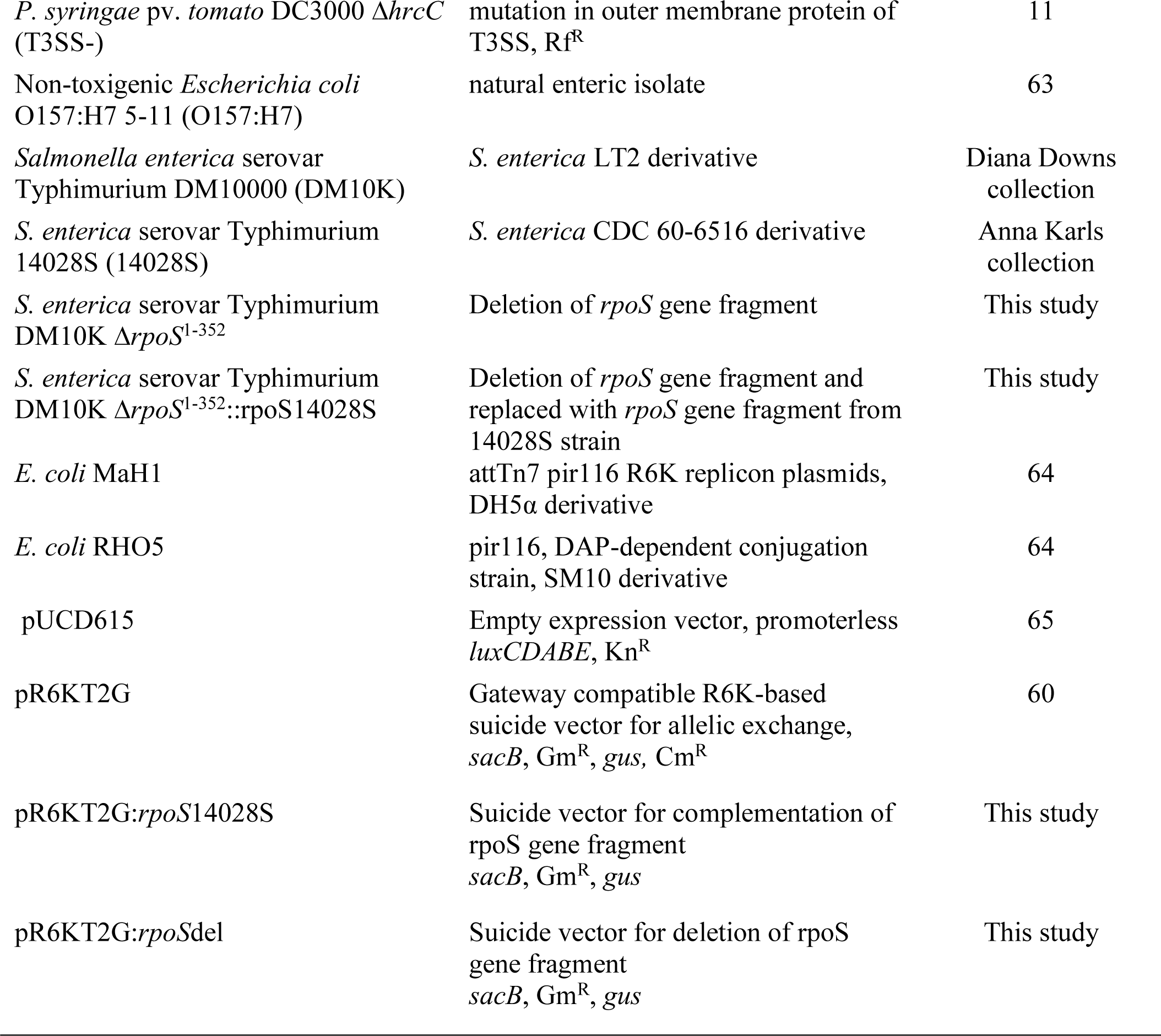
Strains and plasmids used in this study.

First, a set of inocula with a consistent initial concentration of O157:H7 (5 x10^4^ CFU mL^-1^ for *A. thaliana* and 5 x 10^5^ CFU mL^-1^ for *N. benthamiana*) and varying initial DC3K concentrations were infiltrated into both model hosts. Comparative analyses of bacterial colonization at 3 days post inoculation (dpi) of DC3K strains revealed that the T3SS+ strain reached significantly higher bacterial loads than the T3SS-strain in *A. thaliana* regardless of the initial DC3K population (Fig 1A). This suggests that successful infection by the compatible pathogen occurred by suppressing PTI. In all inocula tested where the DC3K initial population varied, DC3K promoted the growth of O157:H7 as plant disease was established in *A. thaliana* regardless of the initial DC3K population (Fig 1A). This suggests that, disease establishment can create a permissive environment in the apoplast for opportunistic colonization of *A. thaliana* by O157:H7.

**Fig 1.**
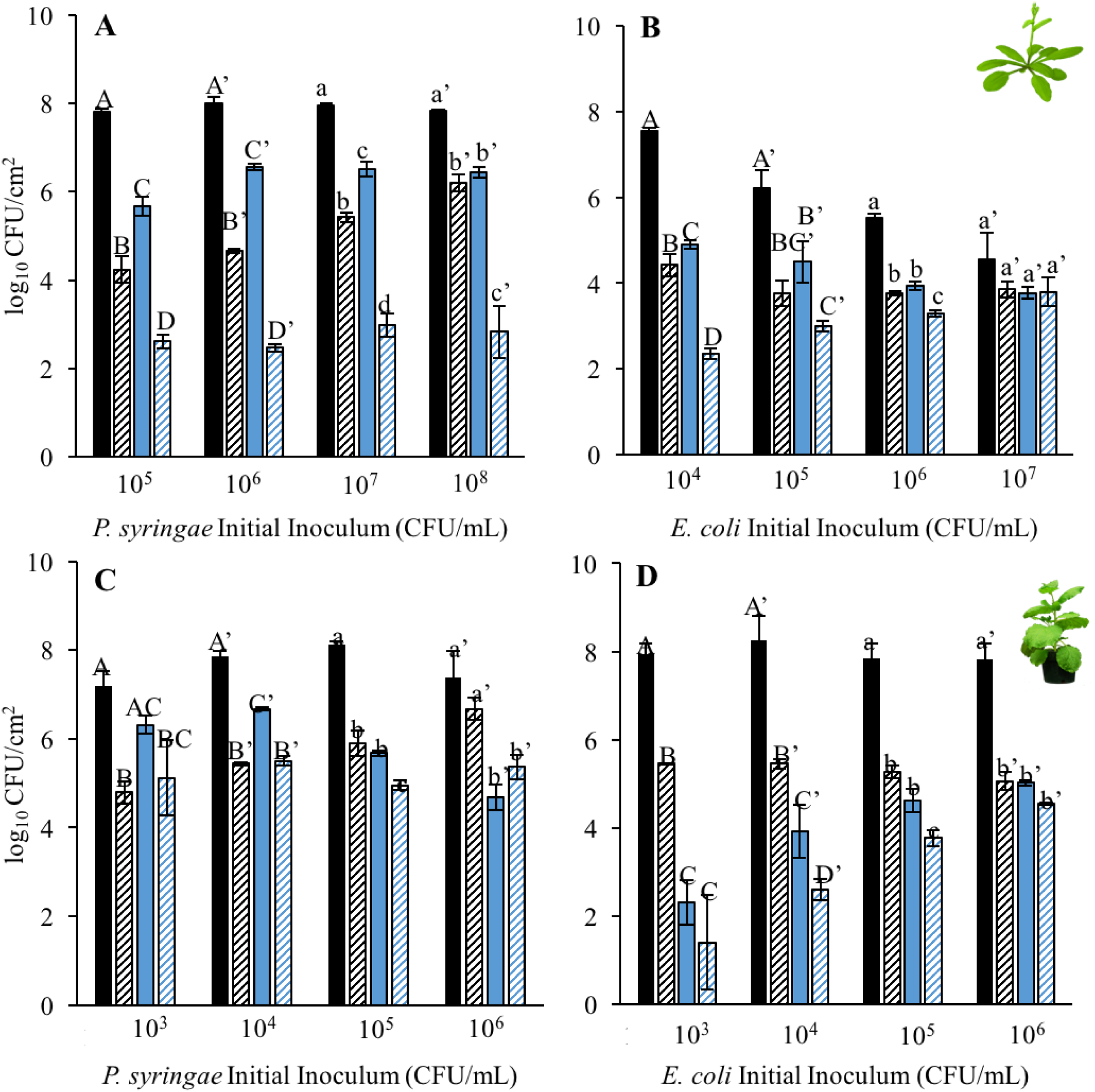
Increased endophytic colonization of *E. coli* O157:H7 during plant disease established by *P. syringae* is dependent on both *E. coli* and *P. syringae* initial populations. Bacterial populations of *P. syringae* pv. *tomato* DC3000 or DC3000Δ*hopQ1-1* with (black) and without (striped) a functional Type III Secretion System and co-inoculation partner, non-toxigenic *E. coli* O157:H7 5-11 (blue) were suspended in 0.25 mM MgCl_2_ and inoculated into *A. thaliana* Col-0 and *N. benthamiana* leaves. **AC)** *E. coli* was inoculated into leaves at a concentration of 5 x 10^4-5^ CFU mL^-1^ with varying concentrations of *P. syringae*. **BD**) *P. syringae* was inoculated into leaves at a concentration of 5 x 10^4-5^ CFU mL^-1^ with varying concentrations of *E. coli*. Bacterial populations were measured as log colony forming units per cm^2^ of leaf tissue (log_10_ CFU/cm^2^) 3 days post-inoculation. Data are means ± SD (n = 3 plants). Different letters indicate significant differences (1-way ANOVA for each inoculum density at p < 0.05).

In contrast, there was no difference in colonization of T3SS+ and T3SS-strains in *N. benthamiana* with the highest (5 x 10^6^ CFU mL^-1^) initial DC3K populations, and there was no consistent difference in O157:H7 colonization regardless of co-inoculation partner (Fig 1C), although a moderate increase in O157:H7 was observed with the T3SS+ strain when this was inoculated at 5 x 10^4^ or 5 x 10^5^ CFU mL^-1^.

Using this information, a set of inocula with a consistent initial concentration of DC3K (5 x10^5^ CFU mL^-1^ for *A. thaliana* and 5 x 10^4^ CFU mL^-1^ for *N. benthamiana*) and varying initial O157:H7 concentrations were infiltrated into both model hosts. We demonstrate that the initial concentration of O157:H7 can influence their opportunistic growth during plant disease. With increasing O157:H7 populations in *A. thaliana*, there was a corresponding decrease in T3SS+ DC3K populations, and with the highest (5 x 10^7^ CFU mL^-1^) O157:H7 initial population, disease was unable to be established by DC3K resulting in no difference in O157:H7 colonization regardless of its co-inoculation partner. (Fig 1B). This suggests that the ratio of plant pathogen to human enteric pathogen strain can affect the ability of DC3K to suppress host immunity or to colonize host tissues.

In *N. benthamiana*, disease was established in all conditions, and high (5 x 10^6^ CFU mL^-^ and low (5 x 10^3^ CFU mL^-1^) O157:H7 initial populations resulted in no difference in colonization by DC3KΔ*hop*Q1-1 regardless of co-inoculation partner (Fig 1D). As in *A. thaliana*, an initial inoculum of 100 times more O157:H7 than DC3K in *N. benthamiana* resulted in a non-permissive environment for opportunistic growth of O157:H7 that was independent of disease establishment. Overall, the *N. benthamiana* apoplast was more restrictive for enhanced opportunistic colonization of O157:H7 than in *A. thaliana*. Based on our findings, the ability for human enteric pathogens to opportunistically colonize plant tissue during plant disease is dependent on the initial populations of both the plant pathogen and human enteric pathogen and on the host species.

### Increased endophytic colonization of *S. enterica* during plant disease established by *P. syringae* is dependent on both host and strain factors

We expanded our study of opportunistic colonization of human enteric pathogens during plant disease in our two model hosts by including additional human enteric pathogen strains. Co-inoculation assays were performed on adult *A. thaliana* and *N. benthamiana* plants by syringe infiltration using two compatible DC3K strains (DC3K and DC3KΔ*hopQ1-1*) with a functional T3SS (T3SS+) or a DC3K strain without a functional T3SS (DC3KΔ*hrcC*, T3SS-) and one of three human enteric pathogen strains, *S. enterica* serovar Typhimurium DM10000 (DM10K), *S. enterica* serovar Typhimurium 14028S (14028S), and *E. coli* O157:H7 (Table 1). These human enteric strains were selected for this study as they are non-pathogenic and have well annotated genomes. *S. enterica* and *E. coli* bacterial suspensions were also infiltrated into plants without a co-inoculation partner as a control. We used the initial inoculum range from above-mentioned experimental results and previous published co-inoculation methods (23) to inform our decision on the initial inoculum concentrations of *P. syringae*, *E. coli* and *S. enterica* to use in our subsequent co-inoculation study in order to ensure that a permissive environment was generated in each host apoplast during plant disease. In *A. thaliana*, human enteric pathogen initial populations never exceeded that of DC3K. In *N. benthamiana,* DC3K initial populations did not exceed 5 x 10^5^ CFU mL^-1^ and human enteric pathogen initial populations did not fall below 5 x 10^4^ CFU mL^-1^. In both hosts, DM10K and 14028S had the same initial populations during co-inoculation.

Comparative analyses of bacterial colonization at 3 dpi of DC3K strains revealed that the T3SS+ strains reached significantly higher bacterial loads in both model hosts than the T3SS-strain, suggesting that successful infection by the compatible pathogen occurred by suppressing PTI, as observed previously (Fig S1, Fig 2). Both O157:H7 and 14028S showed significantly greater colonization in both model hosts when co-inoculated with T3SS+ strains than with T3SS-or by itself (Fig S1, Fig 2A, 2C). Therefore, association with the compatible plant pathogen, DC3K or DC3KΔ*hopQ1-1*, can promote the growth of O157:H7 and 14028S and this is most likely due to the suppression of PTI by DC3K effectors delivered by the T3SS. In contrast, DM10K had significantly greater colonization when co-inoculated with T3SS+ than T3SS- or by itself in *A. thaliana* but not in *N. benthamiana* (Fig 2B, 2D). The inability of DM10K to colonize infected *N. benthamiana* leaves suggests that there are host factors that contribute to opportunistic colonization of *S. enterica* during plant disease. Additionally, 14028S exhibited increased growth during plant disease on the same host, *N. benthamiana*, which suggests that there are also strain factors that also contribute to opportunistic colonization of *S. enterica* during plant disease (Figure 2C).

**Fig 2.**
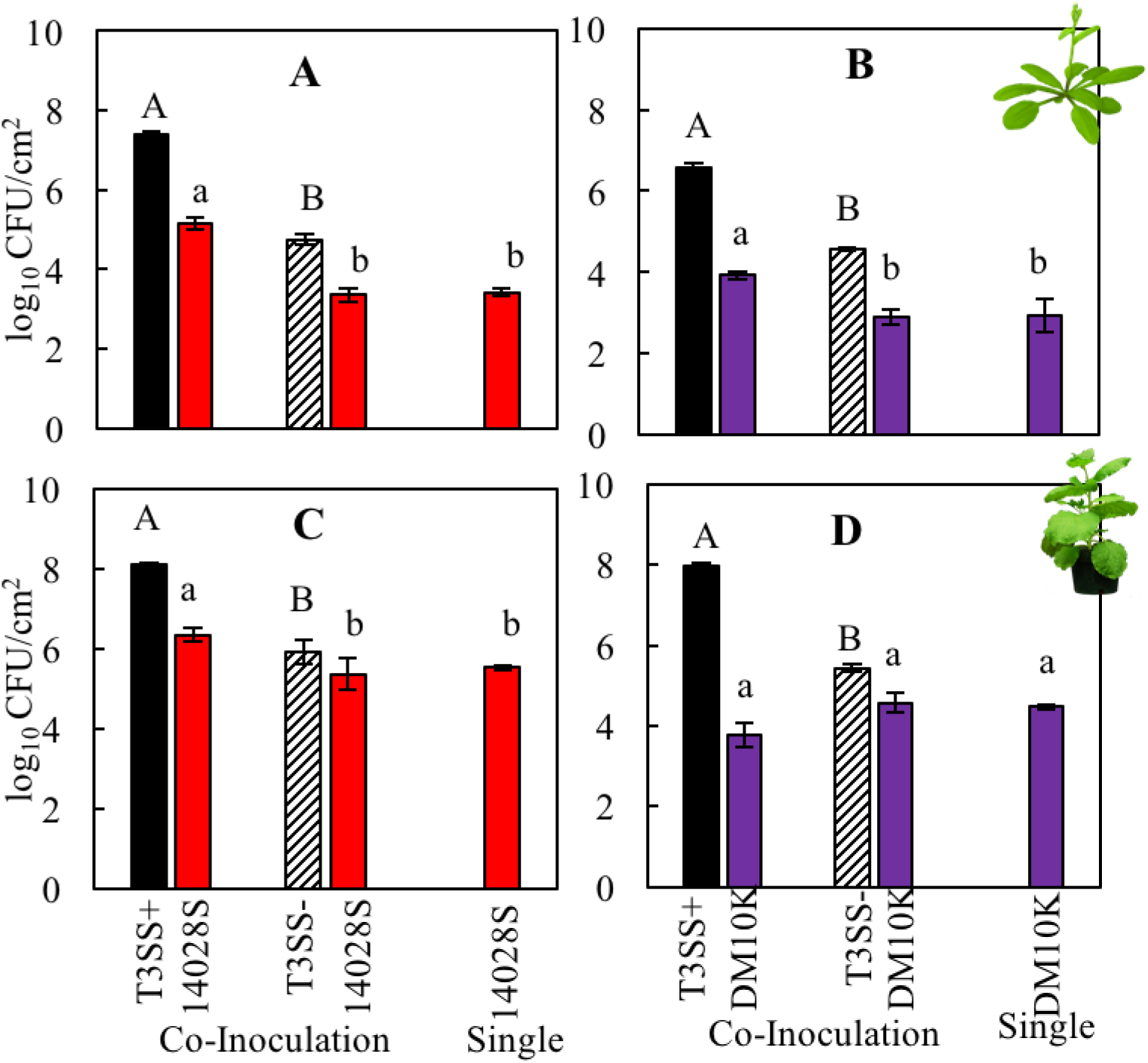
Increased endophytic colonization of *S. enterica* during plant disease established by *P. syringae* is dependent on both host and strain factors. Bacterial populations of *S. enterica* strains **AC**) DM10000 (DM10K, purple), and **BD**) 14028S (14028S, red) with co-inoculation partner *P. syringae* pv. *tomato* DC3000 or DC3000Δ*hopQ1-1* with (T3SS+, black) and without (T3SS-, striped) a functional Type III Secretion System. Inocula were syringe infiltrated into model plant hosts, **AB**) *A. thaliana* Col-0 at a concentration of 5 x 10^6^ CFU mL^-1^ for DC3000 strains and 5 x 10^5^ CFU mL^-1^ for *S. enterica* strains and **CD**) *N. benthamiana* at a concentration of 5 x 10^5^ CFU mL^-1^ for all strains. Bacterial populations were measured as log colony forming units per cm^2^ of leaf tissue (log_10_ CFU/cm^2^) 3 days post-inoculation. Data are means ± SD (n = 3 plants). Different letters indicate significant differences (2-tailed t-test for each strain at p < 0.05).

With foodborne illness outbreaks associated with fresh produce on the rise, we moved our pathosystem from a model host into a crop host. Collards/Kale (*Brassica oleracea* var. *acephala*) and *Arabidopsis* belong the same order, Brassicales, and other *Brassica oleracea* have been previously demonstrated to be compatible hosts for DC3K (27). We demonstrate that DC3K can infect collard leaves in a T3SS dependent manner and develop classic bacterial spot symptoms after syringe inoculation on adult leaves and after spray inoculation on seedlings (Fig S2). As in *A. thaliana*, all human enteric pathogen strains had significantly greater colonization in collards when co-inoculated with T3SS+ than with T3SS- or by itself (Fig 3). Therefore, diseased crops could serve as a potential source for endophytic colonization of human enteric pathogens such as *S. enterica* and *E. coli*.

**Fig 3.**
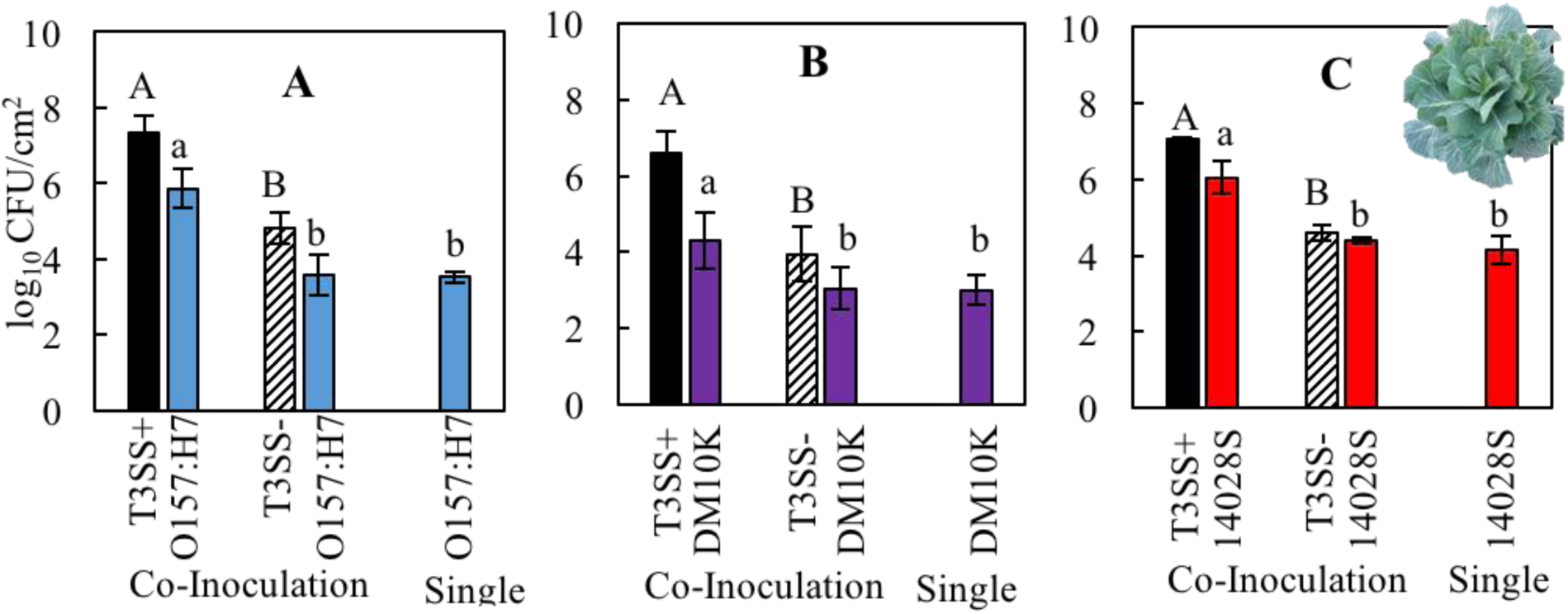
Increased endophytic colonization of *S. enterica* and *E. coli* strains during plant disease established by *P. syringae* in collards. Bacterial populations of human enteric pathogens **A**) non-toxigenic *E. coli* O157:H7 5-11 (O157:H7, blue), **B**) *S. enterica* DM10000 (DM10K, purple), and **C**) *S. enterica* 14028S (14028S, red) with co-inoculation partner *P. syringae* pv. *tomato* DC3000 with (T3SS+, black) and without (T3SS-, striped) a functional Type III Secretion System. Inocula were syringe infiltrated into *B. oleracea* var. *acephala* at a concentration of 5 x 10^5^ CFU mL^-1^ for all strains. Bacterial populations were measured as log colony forming units per cm^2^ of leaf tissue (log_10_ CFU/cm^2^) 3 days post-inoculation. Data are means ± SD (n = 3 plants). Different letters indicate significant differences (2-tailed t-test for each strain at p < 0.05).

### *P. syringae* exhibits reduced growth in *Nicotiana benthamiana* apoplastic wash fluid in the presence of *S. enterica*

To further explore the strain factors that contribute to the opportunistic endophytic colonization of *S. enterica* during plant disease, we posed three hypotheses that may explain why DM10K is unable to colonize *N. benthamiana* during disease by DC3K: 1) During co-inoculation DC3K outcompetes DM10K for limited resources in the apoplast. 2) The two strains, DM10K and 14028S, differ in their ability to metabolize nutrients within the *N. benthamiana* apoplast. 3) There are factors constitutively present or induced during infection to which DM10K is more sensitive than 14028S.

To determine if DC3K outcompetes DM10K for shared resources in the apoplast, we grew DM10K and 14028S as well as DC3K together and separately in M9 minimal media and in *N. benthamiana* apoplastic wash fluid (BAWF) extracted from *N. benthamiana* leaves. Their initial and final populations were measured by dilution plating after peak growth was achieved. Peak growth, measured as maximum OD_600_, was determined by analyzing growth curves in each media type and was determined to be 24 h for BAWF and 48 h for M9 minimal media (Fig S3). Final populations of *S. enterica* were significantly greater than that of DC3K when grown separately in M9 minimal media (Fig S3A, Fig 4A). Similarly, after co-inoculation with DC3K, both DM10K and 14028S strains grew to a significantly higher final population compared to their DC3K co-inoculation partner (Fig 4A). Although initial 14028S populations were four times greater than that of its co-inoculation partner, final 14028S populations were 16 times greater than DC3K. Therefore, the observed increase in 14028S population over that of DC3K is likely not a direct result of its higher initial population. The observed differences in growth between *S. enterica* strains and DC3K in minimal media may be due to differences in doubling time between these two species. Single populations of both *S. enterica* strains grew to an equivalent or significantly higher population alone than when co-inoculated with DC3K in M9 minimal media (Fig 4A). In contrast, final DC3K populations were significantly higher when co-inoculated with an *S. enterica* partner than alone. However, these differences were maintained from higher initial DC3K populations during co-inoculation (Fig 4A). This suggests, that DC3K does not benefit from co-inoculation with any *S. enterica* strain.

**Fig 4.**
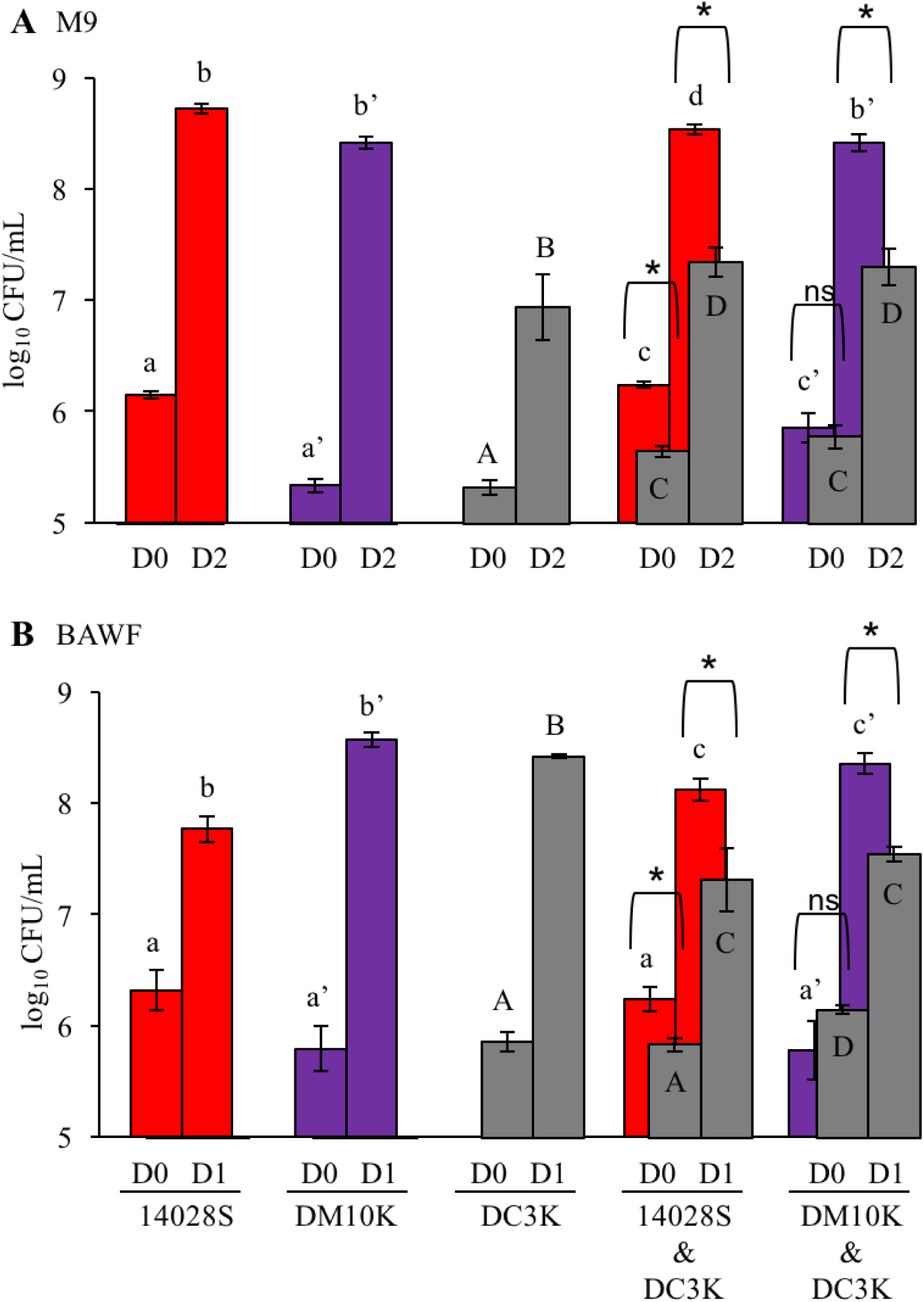
*P. syringae* exhibits reduced growth in *Nicotiana benthamiana* apoplastic wash fluid in the presence of *S. enterica*. Bacterial populations of *P. syringae* pv. *tomato* DC3000 (DC3K, grey) and *S. enterica* strains DM10000 (DM10K, purple) and 14028S (red) after inoculation in **A**) M9 minimal media and **B**) *N. benthamiana* apoplastic wash fluid (BAWF). Bacterial populations were measured as log colony forming units per mL of culture (log_10_ CFU/mL) on day 0, 1, and 2. Data are means ± SD (n = 3-5). Different letters indicate significant differences (2-way ANOVA for each strain at p < 0.05). Asterisk indicates significant difference between co-inoculation partners (2 tailed t-test at p < 0.05).

In BAWF, both DM10K and 14028S strains grew to a significantly higher final population compared to their DC3K co-inoculation partner (Fig 4B). Additionally, 14028S initial populations were 3 times greater than that of DC3K and 7 times greater after 1 day of growth in BAWF. Therefore, the observed increase in 14028S population over that of DC3K is likely not a direct result of its higher initial population. Both DC3K and DM10K strains grew to a significantly higher population alone than when co-inoculated together. In contrast, 14028S had a significantly less population alone than when co-inoculated with DC3K despite having similar initial populations (Fig 4B). This suggests that 14028S benefits from DC3K co-inoculation in BAWF whereas DM10K does not.

### *S. enterica* strain DM10000 exhibits a pronounced biphasic growth pattern in *N. benthamiana* apoplastic wash fluid that is suppressed by exogenous glucose and phosphate

A second potential explanation as to why the DM10K is unable to colonize *N. benthamiana* during disease by DC3K is that the two *S. enterica* strains differ in their ability to metabolize host-derived nutrients. To test this, we grew DM10K and 14028S in rich media (Luria Broth (LB)), minimal media (M9), and apoplastic wash fluid collected from the two model hosts, *A. thaliana* and *N. benthamiana*. Both strains have similar growth in rich media and minimal media which suggests that the two strains have similar metabolic potentials (Fig 5A&B). In apoplastic wash fluid, DM10K grows to a higher density than 14028S with biphasic growth in BAWF, whereas both strains have similar growth in *A. thaliana* apoplastic wash fluid (Fig 5C&D). This suggests that the two *S. enterica* strains differ in their ability to metabolize one or more specific *N. benthamiana-*derived nutrients The biphasic growth pattern may be indicative of two different nutrient sources in BAWF being preferentially metabolized by *S. enterica* which implies differential metabolism of BAWF nutrients by these two strains.

**Fig 5.**
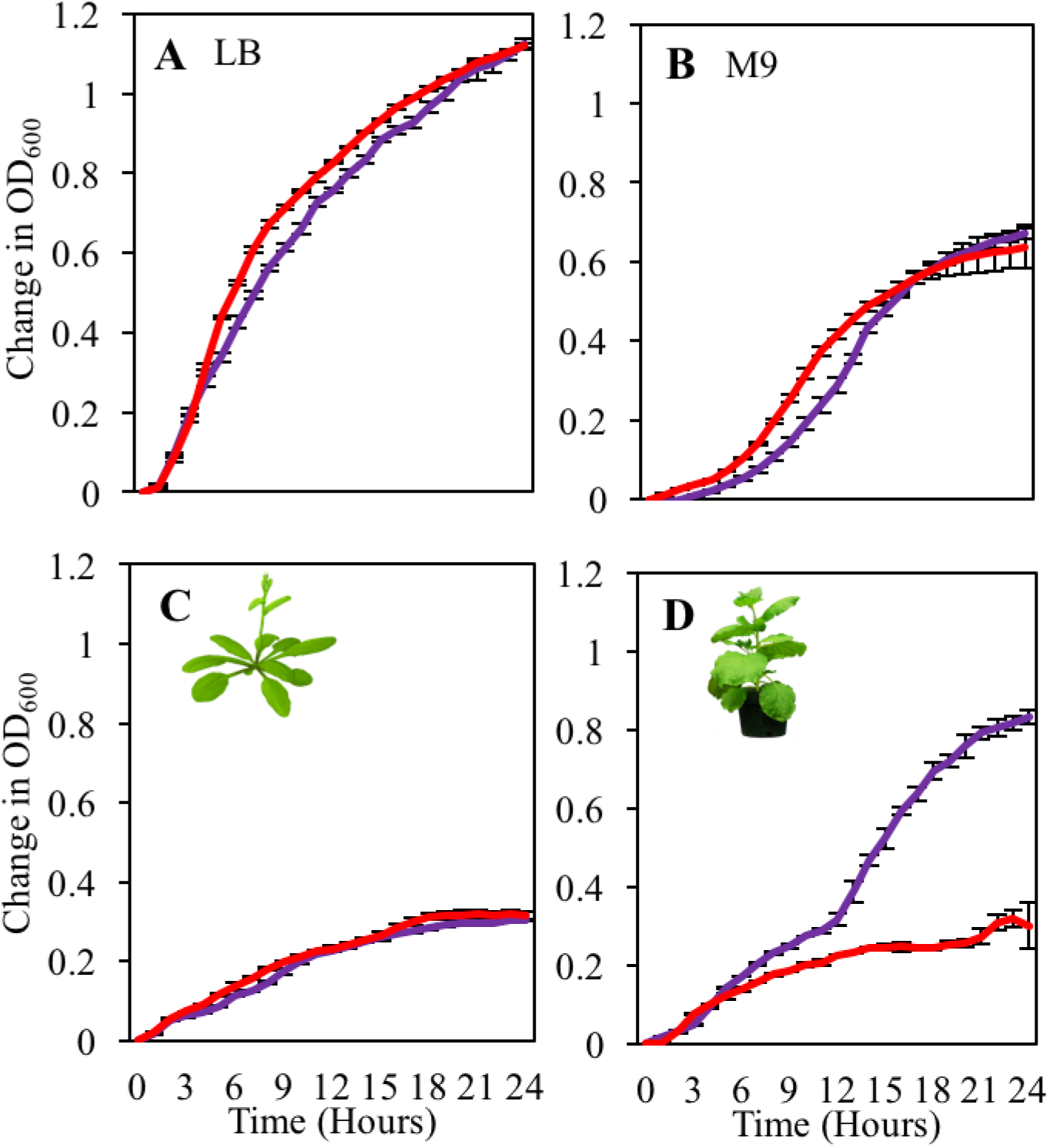
*S. enterica* strain DM10000 exhibits greater growth and a biphasic growth pattern in *Nicotiana benthamiana* apoplastic wash fluid. Growth curves of *S. enterica* DM10000 (purple) and *S. enterica* 14028S (red) in **A**) Luria Broth (LB) **B**) M9 minimal media, **C**) filtered *A. thaliana* apoplastic wash fluid, and **D**) filtered *N. benthamiana* apoplastic wash fluid. Cultures were incubated at 22°C, and the OD_600_ was recorded every hour. Growth was measured as the mean change in OD_600_. Error bars show standard deviation (n=5 wells).

As this biphasic growth response could be linked to catabolite repression of apoplastic derived nutrient utilization, we aimed to test what compounds alter this biphasic growth pattern. Both *S. enterica* strains were grown in BAWF supplemented with exogenous macronutrients and micronutrients using water as a control. The concentrations of the macronutrients and micronutrients were determined based on concentrations found in the M9 minimal media. The DM10K strain grew to a higher population than the 14028S strain with both exhibiting biphasic growth in BAWF supplemented with sodium chloride, magnesium sulfate, ammonium sulfate, and calcium chloride similarly to that of the water control (Fig S4). We found that *S. enterica* biphasic growth in BAWF was suppressed when supplemented with exogenous glucose and potassium phosphate compared to the water control (Fig 6). DM10K biphasic growth was more starkly suppressed by these two compounds compared to 14028S. Additionally, supplementation with potassium phosphate had a temporary effect on biphasic growth suppression compared to glucose (Fig 6). We confirmed that it was the phosphate anion and not the potassium cation that suppressed biphasic growth by supplementing BAWF with potassium chloride, potassium phosphate, and sodium phosphate using water as a control. In both cases where a phosphate anion was supplemented, *S. enterica* biphasic growth was suppressed and supplementation with potassium chloride resulted in similar biphasic growth to that of the water control (Fig S5). The suppression of biphasic growth of the two *S. enterica* strains by glucose and phosphate in BAWF is indicative of a catabolite repression response. Glucose is a preferred carbon source that is likely metabolized by the *S. enterica* strains first which represses the enzyme system required for the metabolism of other carbon sources available in BAWF. A better understanding of what carbon sources within BAWF are differentially metabolized by these two *S. enterica* strains may provide insight into their differential colonization during plant disease in *N. benthamiana*.

**Fig 6.**
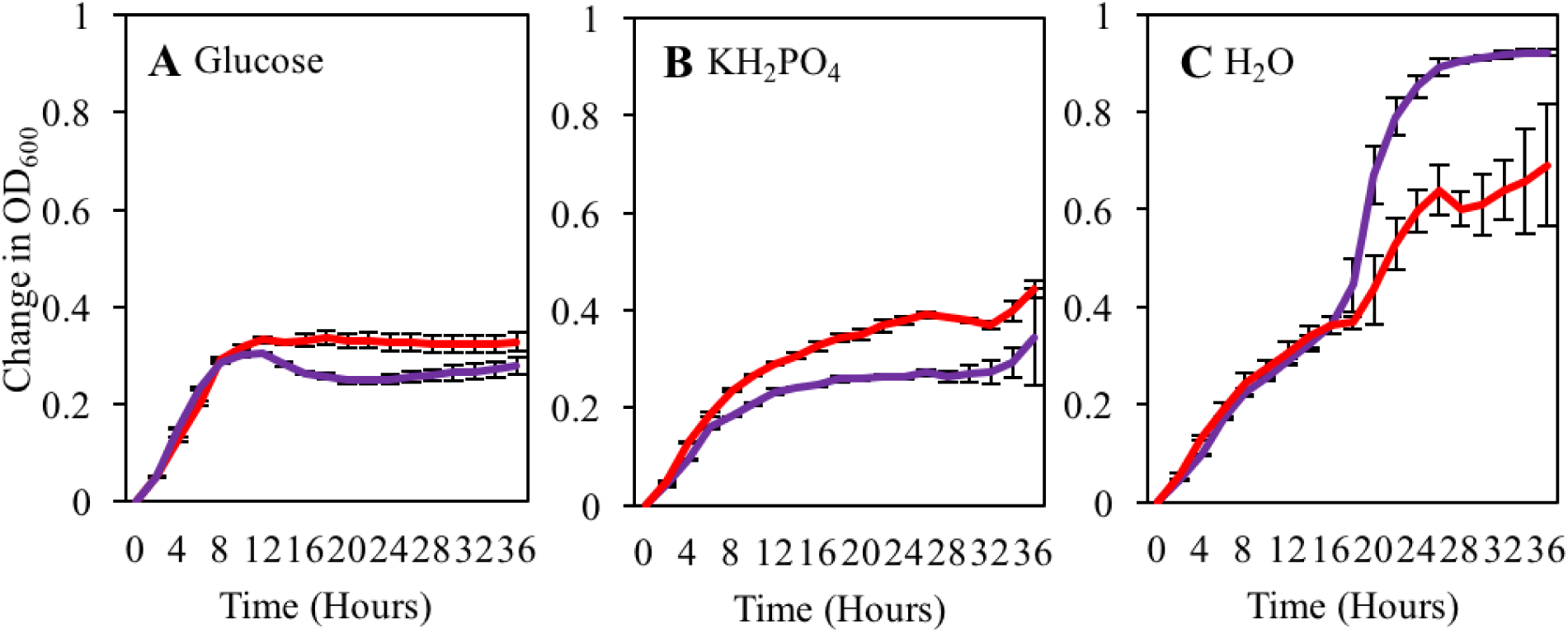
*S. enterica* strain biphasic growth in *Nicotiana benthamiana* apoplastic wash fluid is suppressed by exogenous glucose and phosphate. Growth curves of *S. enterica* DM10000 (purple) and *S. enterica* 14028S (red) in filtered *Nicotiana benthamiana* apoplastic wash fluid supplemented with **A**) 25 mM glucose, **B**) 10 mM potassium phosphate, and **C**) water. Cultures were aliquoted into 5 replicate wells, incubated at 22°C, and the OD_600_ was recorded every hour. Growth was measured as the average change in OD_600_. Error bars show standard deviation (n=5 wells).

### *S. enterica* strain 14028S metabolizes a more diverse range of apoplastic derived carbon sources than *S. enterica* strain DM10000

The differential growth response between *S. enterica* DM10K and *S. enterica* 14028S in in vitro growth assays could be due to difference in the ability of these bacteria to use carbon metabolites present in BAWF. We used the Biolog Phenotypic MicroArray™ system to generate carbon utilization profiles for our two strains as described by Rico and Preston (28). Bacterial cultures were exposed to BAWF or rich media (LB) for three hours, after which cultures were inoculated on PM1 MicroPlates™ and incubated for 24 hours. These plates contain 95 unique carbon sources and the metabolic indicator tetrazolium violet. Reduction of the tetrazolium violet is indicative that the strain is able to metabolize that specific carbon source. This gave an indication of the metabolic potential of these two strains when exposed to our two media types.

An overview of the carbon utilization results for our *S. enterica* strains is provided in Table 2. Both DM10K and 14028S use a common set of 80 substrates as carbon sources regardless of which media they were pre-cultured in. These 80 substrates include 18 sugar or sugar derivatives, 3 sugar alcohols, 23 organic acids, and 17 amino acids or peptides. However, 12 carbon sources showed variable distribution between strains and/or media types. More specifically, the DM10K strain could metabolize 6 unique carbon sources when previously pre-cultured BAWF but not LB including sucrose, α-D-lactose, lactulose, D-cellobiose, D-malic acid, and adonitol. In contrast, the DM10K strain could only metabolize formic acid as a carbon source when exposed to LB but not BAWF. For the 14028S strain, 3 unique carbon sources were metabolized when exposed to BAWF but not LB including α-hydroxy glutaric acid-ɣ-lactone, glycolic acid, and 2-aminoethanol. The 14028S strain could only metabolize acetoacetic acid as a carbon source when exposed to LB but not BAWF.

**Table 2.** Carbon source utilization by S. enterica strains DM10000 and 14028S in N. benthamiana apoplastic wash fluid and rich media.

| Carbon Source <sup>b</sup> | Un-inhibited Signal <sup>a</sup> |  |  |  | Inhibited Signal <sup>a</sup> |  |  |  |
| --- | --- | --- | --- | --- | --- | --- | --- | --- |
|  | DM10K |  | 14028S |  | DM10K |  | 14028S |  |
|  | LB | BAWF | LB | BAWF | LB | BAWF | LB | BAWF |
| p-Hydroxy Phenyl Acetic Acid | 1.23 | 1.36 | 1.06 | 0.99 | 0.03 | -0.02 | 0.11 | -0.01 |
| m-Hydroxy Phenyl Acetic Acid | 1.21 | 1.36 | 1.10 | 1.03 | 0.03 | -0.04 | 0.17 | 0.03 |
| Mucic Acid * | 1.30 | 1.32 | 1.05 | 0.91 | 0.09 | -0.02 | 0.03 | 0.10 |
| N-Acetyl-β-D-Mannosamine * | 1.38 | 1.30 | 1.16 | 0.98 | 0.23 | -0.01 | 0.29 | 0.27 |
| Tyramine | 1.33 | 1.28 | 1.12 | 1.06 | 0.04 | -0.04 | 0.03 | 0.01 |
| Maltotriose *# | 1.28 | 1.26 | 1.06 | 0.83 | 0.11 | 0.07 | 0.21 | 0.26 |
| Fumaric Acid * | 1.35 | 1.26 | 1.00 | 0.85 | 0.26 | 0.01 | 0.20 | 0.33 |
| Inosine * | 1.17 | 1.23 | 1.16 | 1.03 | 0.28 | 0.04 | 0.48 | 0.09 |
| L-Malic Acid * | 1.15 | 1.22 | 0.99 | 0.97 | 0.39 | -0.03 | 0.14 | 0.28 |
| D-Glucosiminic Acid | 1.31 | 1.20 | 1.00 | 0.99 | 0.04 | -0.02 | 0.29 | 0.01 |
| D-Gluconic Acid * | 1.33 | 1.19 | 1.15 | 1.12 | 0.17 | 0.02 | 0.19 | 0.23 |
| Tricarballic Acid | 1.28 | 1.16 | 1.03 | 0.97 | 0.02 | 0.03 | 0.27 | 0.02 |
| D-Glucuronic Acid | 1.30 | 1.14 | 1.14 | 1.04 | 0.08 | 0.05 | 0.34 | 0.04 |
| L-Alanine * | 1.26 | 1.13 | 0.93 | 0.87 | 0.08 | 0.05 | 0.16 | 0.05 |
| L-Serine * | 1.30 | 1.12 | 1.09 | 0.93 | 0.30 | -0.01 | 0.38 | 0.10 |
| L-Lactic Acid * | 1.26 | 1.11 | 1.07 | 0.94 | 0.21 | -0.01 | 0.23 | 0.31 |
| D-Glucose-1-Phosphate * | 1.30 | 1.08 | 1.03 | 1.01 | 0.26 | 0.04 | 0.11 | 0.34 |
| D-Fructose-6-Phosphate * | 1.28 | 1.06 | 0.98 | 0.95 | 0.24 | -0.04 | 0.07 | 0.27 |
| D,L-Malic Acid * | 1.30 | 1.05 | 0.99 | 0.93 | 0.40 | -0.01 | 0.11 | 0.30 |
| L-Proline | 1.22 | 1.05 | 1.22 | 1.11 | 0.07 | 0.00 | 0.12 | 0.02 |
| Pyruvic Acid * | 1.13 | 1.04 | 0.99 | 0.92 | 0.19 | -0.01 | 0.32 | 0.19 |
| Uridine | 1.15 | 1.03 | 1.08 | 1.01 | 0.24 | -0.01 | 0.07 | 0.04 |
| Glycyl-L-Proline * | 1.40 | 1.02 | 0.68 | 0.62 | 0.01 | -0.05 | -0.02 | 0.08 |
| Propionic Acid * | 1.12 | 1.02 | 0.90 | 0.71 | 0.11 | -0.01 | 0.03 | 0.16 |
| Citric Acid | 1.23 | 1.02 | 0.93 | 0.93 | 0.04 | 0.00 | 0.01 | -0.07 |
| D-Melibiose | 1.14 | 1.01 | 0.94 | 0.67 | 0.03 | -0.02 | 0.02 | -0.05 |
| α-Methyl-D-Galactoside | 1.31 | 1.01 | 1.02 | 0.78 | 0.04 | -0.01 | -0.03 | -0.05 |
| Glycerol * | 1.18 | 1.00 | 0.94 | 0.80 | 0.14 | 0.01 | 0.35 | 0.20 |
| L-Aspartic Acid *# | 0.99 | 0.98 | 1.08 | 0.98 | 0.30 | 0.06 | 0.20 | 0.34 |
| L-Asparagine * | 1.14 | 0.95 | 0.85 | 1.06 | 0.24 | 0.02 | 0.21 | 0.12 |
| Succinic Acid * | 0.89 | 0.92 | 0.94 | 0.89 | 0.27 | 0.00 | 0.43 | 0.30 |
| 2-Deoxy Adenosine * | 1.11 | 0.92 | 1.08 | 0.96 | 0.09 | -0.02 | 0.23 | 0.34 |
| D-Serine | 1.30 | 0.90 | 0.86 | 1.01 | 0.00 | -0.02 | 0.16 | 0.04 |
| D-Saccharic Acid | 1.16 | 0.89 | 0.98 | 0.95 | 0.05 | -0.01 | 0.04 | -0.05 |
| Dulcitol | 1.07 | 0.89 | 0.43 | 0.25 | 0.03 | -0.02 | -0.04 | -0.03 |
| D-Glucose-6-Phosphate * | 1.10 | 0.89 | 0.90 | 0.97 | 0.11 | 0.02 | 0.29 | 0.27 |
| Adenosine * | 1.02 | 0.88 | 0.85 | 0.80 | 0.22 | 0.00 | 0.13 | 0.38 |
| L-Alanyl-Glycine * | 1.31 | 0.88 | 1.01 | 0.95 | 0.10 | 0.01 | 0.15 | 0.14 |
| D-Xylose | 1.07 | 0.86 | 0.64 | 0.30 | -0.04 | -0.10 | 0.05 | -0.10 |
| Thymidine * | 1.04 | 0.86 | 1.06 | 0.93 | 0.08 | 0.00 | 0.11 | 0.21 |
| S-Sorbitol | 1.14 | 0.85 | 0.69 | 0.66 | 0.05 | 0.00 | 0.06 | -0.06 |
| L-Fucose | 1.17 | 0.85 | 0.79 | 0.60 | 0.03 | 0.00 | 0.16 | -0.04 |
| Methyl Pyruvate * | 0.80 | 0.84 | 0.81 | 0.79 | 0.26 | 0.00 | 0.22 | 0.14 |
| D-Fructose * | 1.04 | 0.83 | 0.67 | 0.45 | 0.36 | 0.04 | 0.24 | 0.23 |
| D-Galactonic Acid-γ-Lactone | 1.32 | 0.81 | 1.07 | 0.46 | 0.01 | 0.02 | 0.02 | -0.05 |
| α-D-Glucose * | 1.01 | 0.81 | 0.59 | 0.36 | 0.34 | 0.03 | 0.15 | 0.17 |
| D-Mannitol * | 0.97 | 0.80 | 0.57 | 0.25 | 0.23 | 0.00 | 0.14 | 0.19 |
| D,L-α-Glycerol-Phosphate * | 0.84 | 0.80 | 0.96 | 0.86 | 0.21 | -0.01 | 0.35 | 0.08 |
| Glycyl-L-Glutamic Acid * | 1.36 | 0.79 | 0.34 | 0.32 | 0.04 | -0.02 | 0.19 | 0.12 |
| D-Alanine | 0.80 | 0.79 | 0.82 | 0.53 | 0.01 | -0.04 | 0.01 | -0.12 |
| L-Rhamnose | 1.04 | 0.79 | 0.47 | 0.25 | 0.03 | 0.00 | 0.19 | -0.03 |
| D-Ribose * | 0.93 | 0.78 | 0.56 | 0.57 | 0.17 | -0.13 | 0.10 | 0.09 |
| m-Inositol | 0.93 | 0.76 | 0.51 | 0.76 | 0.07 | 0.00 | 0.09 | -0.06 |
| Acetic Acid * | 0.80 | 0.74 | 0.76 | 0.60 | 0.09 | -0.01 | 0.24 | 0.13 |
| α-Keto-Butyric Acid | 0.57 | 0.74 | 0.66 | 0.47 | 0.01 | 0.02 | 0.17 | -0.04 |
| D-Galactose * | 0.90 | 0.70 | 0.57 | 0.34 | 0.09 | -0.02 | 0.10 | 0.25 |
| α-Hydroxy Butyric Acid * | 0.79 | 0.69 | 0.67 | 0.57 | 0.14 | -0.01 | 0.17 | 0.18 |
| Bromo Succinic Acid * | 0.78 | 0.64 | 0.41 | 0.47 | 0.23 | 0.00 | 0.34 | 0.09 |
| Glycyl-L-Aspartic Acid * | 1.09 | 0.63 | 0.56 | 0.37 | 0.07 | -0.02 | 0.24 | 0.20 |
| D-Mannose * | 0.81 | 0.63 | 0.40 | 0.17 | 0.24 | 0.00 | -0.04 | 0.09 |
| N-Acetyl-D-Glucosamine * | 0.83 | 0.61 | 0.42 | 0.15 | 0.21 | -0.01 | 0.05 | 0.19 |
| Maltose | 0.43 | 0.57 | 0.81 | 0.70 | 0.04 | 0.01 | 0.09 | -0.01 |
| m-Tartaric Acid * | 0.82 | 0.53 | 0.37 | 0.62 | 0.01 | -0.02 | 0.02 | 0.07 |
| D-Trehalose * | 0.68 | 0.51 | 0.52 | 0.32 | 0.05 | -0.01 | 0.06 | 0.14 |
| L-Glutamic Acid * | 0.39 | 0.49 | 0.66 | 0.40 | 0.16 | -0.01 | 0.14 | 0.16 |
| 1,2-Propanediol # | 0.13 | 0.48 | 0.57 | 0.53 | 0.02 | 0.06 | 0.00 | -0.01 |
| Tween 40 | 0.72 | 0.47 | 0.84 | 0.76 | 0.01 | -0.01 | 0.17 | -0.03 |
| L-Arabinose | 0.65 | 0.44 | 0.35 | 0.12 | -0.03 | -0.11 | -0.13 | -0.11 |
| Sucrose | -0.01 | 0.42 | 0.42 | 0.31 | 0.04 | -0.01 | 0.07 | 0.01 |
| α-D-Lactose | 0.02 | 0.41 | 0.43 | 0.38 | 0.03 | 0.02 | 0.21 | -0.05 |
| Tween 20 | 0.76 | 0.41 | 0.84 | 0.76 | 0.01 | -0.02 | 0.09 | -0.02 |
| β-Methyl-D-Glucoside * | 0.35 | 0.40 | 0.64 | 0.48 | 0.15 | -0.01 | 0.05 | 0.23 |
| L-Threonine | 1.09 | 0.38 | 0.28 | 0.09 | 0.13 | 0.01 | 0.22 | 0.05 |
| Tween 80 | 0.46 | 0.36 | 0.66 | 0.64 | 0.01 | -0.02 | 0.01 | -0.02 |
| Glyoxylic Acid | 0.25 | 0.35 | 0.13 | 0.09 | 0.01 | -0.03 | 0.23 | -0.03 |
| D-Aspartic Acid * | 1.00 | 0.30 | 0.83 | 0.61 | 0.17 | 0.02 | 0.32 | 0.11 |
| α-Keto-Glutaric Acid * | 0.12 | 0.28 | 0.26 | 0.25 | 0.04 | 0.00 | 0.25 | 0.06 |
| D-Psicose | 0.15 | 0.25 | 0.13 | 0.09 | 0.10 | 0.00 | 0.13 | 0.02 |
| Lactulose | 0.01 | 0.24 | 0.43 | 0.33 | 0.04 | 0.00 | 0.25 | -0.03 |
| Mono Methyl Succinate * | 0.14 | 0.23 | 0.19 | 0.16 | 0.17 | 0.00 | 0.11 | 0.22 |
| **L-Lyxose** | 0.31 | 0.19 | 0.15 | 0.12 | -0.17 | -0.22 | 0.01 | -0.12 |
| **D-Cellobiose \*** | -0.06 | 0.16 | 0.28 | 0.27 | 0.06 | 0.01 | 0.05 | 0.06 |
| **D-Malic Acid \*** | -0.04 | 0.15 | -0.07 | 0.01 | 0.07 | 0.01 | 0.02 | 0.22 |
| Acetoacetic Acid * | 0.07 | 0.13 | 0.02 | 0.09 | 0.04 | -0.01 | 0.32 | 0.12 |
| **Adonitol** | 0.01 | 0.11 | 0.17 | 0.17 | 0.05 | 0.01 | 0.02 | -0.01 |
| **L-Glutamine \*** | 0.35 | 0.10 | 0.12 | 0.14 | 0.28 | -0.01 | 0.09 | 0.09 |
| α-Hydroxy Glutaric Acid-γ-Lactone | 0.08 | 0.10 | 0.08 | 0.02 | 0.03 | 0.01 | 0.09 | -0.02 |
| Glucuronamide | 0.08 | 0.08 | -0.02 | -0.10 | 0.01 | -0.02 | 0.12 | -0.04 |
| **D-Threonine** | 0.10 | 0.08 | 0.08 | 0.07 | -0.02 | 0.03 | 0.28 | 0.00 |
| Formic Acid * | 0.27 | 0.04 | 0.69 | 0.32 | 0.24 | -0.02 | 0.39 | 0.18 |
| **Glycolic Acid #** | -0.01 | 0.04 | 0.05 | -0.01 | 0.04 | 0.05 | 0.25 | 0.00 |
| L-Galactonic Acid-γ-Lactone | 0.01 | 0.00 | -0.04 | -0.09 | 0.01 | -0.04 | -0.01 | 0.00 |
| Phenylethyl-amine | -0.06 | 0.00 | 0.00 | -0.10 | 0.05 | 0.00 | 0.03 | -0.01 |
| **D-Galacturonic Acid \*** | -0.05 | -0.01 | 0.00 | -0.09 | 0.03 | 0.02 | 0.26 | 0.10 |
| 2-Aminoethanol | -0.10 | -0.01 | 0.09 | -0.10 | 0.04 | -0.03 | -0.05 | -0.03 |

<sup>a</sup> Given values are average change in normalized optical density at 460 nm ( $\Delta OD_{460}$ ) corrected to the negative control; for each genotype, *Salmonella enterica* DM10000 (DM10K) and *S. enterica* 14028S (14028S), and each growth treatment of rich media (LB) or apoplastic wash fluid collected from *N. benthamiana* leaves (BAWF) the value is the mean of two replicates. Dark gray boxes show positive values ( $\Delta OD_{460} > 0.2$ ), light gray boxes show weak positive values ( $0.05 < \Delta OD_{460} < 0.2$ ), and white boxes show negative values ( $\Delta OD_{460} < 0.05$ ).
<sup>b</sup> Carbon Sources on PM1 plate. Bold carbon sources are sources that have been identified in BAWF via GC-MS. Asterisk (\*) indicates the substrate was used as a carbon source by BAWF inhibitor-treated *S. enterica* 14028S. Pound (#) indicates the substrate was used as a carbon source by BAWF inhibitor-treated *S. enterica* DM10K. Carbon sources are color coded based on their classification: amino acids and peptides (blue), sugars and sugar derivatives (red), organic acids (purple), sugar alcohols (yellow), and other (white). Compounds ranked based on uninhibited signal values of DM10K strain after growth in BAWF.

Comparing carbon utilization profiles between *S. enterica* strains after exposure to BAWF, carbon metabolism was less restrictive in DM10K than 14028S in that it could metabolize 3 unique carbon sources including D-malic acid, α-hydroxy glutaric acid-ɣ-lactone, and glucuronamide. In contrast, comparing carbon utilization profiles between *S. enterica* strains after exposure to LB, carbon metabolism was less restrictive in 14028S than DM10K in that it could metabolize 7 unique carbon sources including sucrose, α-D-lactose, lactulose, D-cellobiose, adonitol, glycolic acid, and 2-aminoethanol. Both *S. enterica* strains failed to use L-galactonic acid-ɣ-lactone, phenylethyl-amine, and D-galacturonic acid.

To identify which carbon utilization pathways are active in our two *S. enterica* strains grown in BAWF and rich media, we used previously-described inhibition assays (28). Bacterial cultures exposed to both BAWF and LB were inhibited with tetracycline prior to inoculation on PM1 MicroPlates™. This inhibitory treatment prevents bacteria from adapting to novel carbon sources by inhibiting protein synthesis. An overview of the inhibitory carbon utilization results is shown in Table 2. Generally, inhibitor-treated strains had more restrictive carbon utilization profiles than uninhibited strains with overall weaker signals corresponding to reduced growth from tetracycline treatment. For the DM10K strain, only 2 carbon sources (L-aspartic acid and maltotriose) were constitutively metabolized regardless of which media it was exposed to. Additionally, DM10K induced the metabolism of 2 carbon sources after exposure to BAWF (1,2-propanediol and glycolic acid) and induced the metabolism of 51 carbon sources after exposure to LB. In contrast, 14028S constitutively metabolized 45 carbon sources regardless of which media it was exposed to. The 14028S strain metabolized 7 carbon sources after exposure to BAWF including mucic acid, glycyl-L-proline, propionic acid, m-tartaric acid, D-mannose, D-malic acid, and β-methyl-D-glucoside, and 26 carbon sources after exposure to LB.

GC-MS analysis of BAWF detected 70 different compounds including, 15 sugar or sugar derivatives, 32 amino acids or amino acid derivatives and 19 organic acids (Table S2). Of the 15 identified BAWF-derived sugar and sugar derivatives, 5 of these were induced as metabolized carbon sources in BAWF inhibitor-treated *S. enterica* strains including D-trehalose, D-glucose, D-fructose, D-galactose, and D-glucose-6-phosphate. Sucrose, myo-inositol, glucose, galactose, and fructose were identified to have the highest relative concentrations in BAWF with values greater than 100 (Table S2). Of the 32 identified BAWF-derived amino acid and amino acid derivatives, 13 of these were induced as metabolized carbon sources in BAWF inhibitor-treated *S. enterica* strains. Proline, alanine, aspartic acid, and pyroglutamic acid were identified to have the highest relative concentrations in BAWF with values greater than 100 (Table S2). Of the 19 BAWF derived organic acids, 8 of these were induced as metabolized carbon sources in BAWF inhibitor-treated *S. enterica* strains including fumaric acid, L-malic acid, D,L-malic acid, succinic acid, glutaric acid, m-tartaric acid, L-lactic acid, and D-malic acid. Malic acid, butanoic acid, quinic acid, and putrescene were identified to have the highest relative concentrations in BAWF with values greater than 100 (Table S2).

### Repairing mutations in the *rpoS* gene of *S. enterica* DM10000 through complementation rescues both *in planta* and *in vitro* phenotypes

As DM10K is an LT2 derivative (Table 1), one of the most notable differences between the LT2 strain and 14028S is the presence of a mutation in the start codon of the *rpoS* sigma factor gene which results in lower RpoS protein levels (29). RpoS (σ38, σ^S^) is a regulatory protein that plays a crucial role in virulence, stress response, and fitness (30, 31). We confirmed through sequencing that our DM10K strain has the expected alternative start codon TTG in the *rpoS* gene as well as an additional 8 bp deletion after the first 114 codons, resulting in a premature stop codon (Figure 7A). To investigate whether these mutations in the *rpoS* gene in DM10K could explain the differences observed in its ability grow in BAWF and inability to colonize during plant disease, we deleted the first 352 bp in the DM10K *rpoS* gene and complemented the DM10KΔrpoS^1-352^ mutant with the 352 bp *rpoS* gene fragment from 14028S through allelic exchange. Confirmed complement strain DM10KΔrpoS^1-352^::rpoS14028S had both large and small colony sizes. The colony sizes and morphology were maintained after sub-culturing onto LB with all strains exhibiting smooth, circular, off-white colonies (Figure S6A). Complement strain DM10KΔrpoS^1-352^::rpoS14028S L1 had significantly smaller colony diameter similar to its parental strain DM10K compared to 14028S and the DM10KΔrpoS^1-352^ deletion strain (Figure S6B). Complement colony B1 had significantly larger colony diameter similar to 14028S compared to its parental strain DM10K (Figure S6B). This suggests that the differences in colony sizes are not due to differences in *rpoS* gene functionality as the DM10KΔrpoS^1-352^ deletion strain containing a dysfunctional *rpoS* gene and 14028S strain containing a fully functional *rpoS* gene have similar colony sizes.

**Fig 7.**
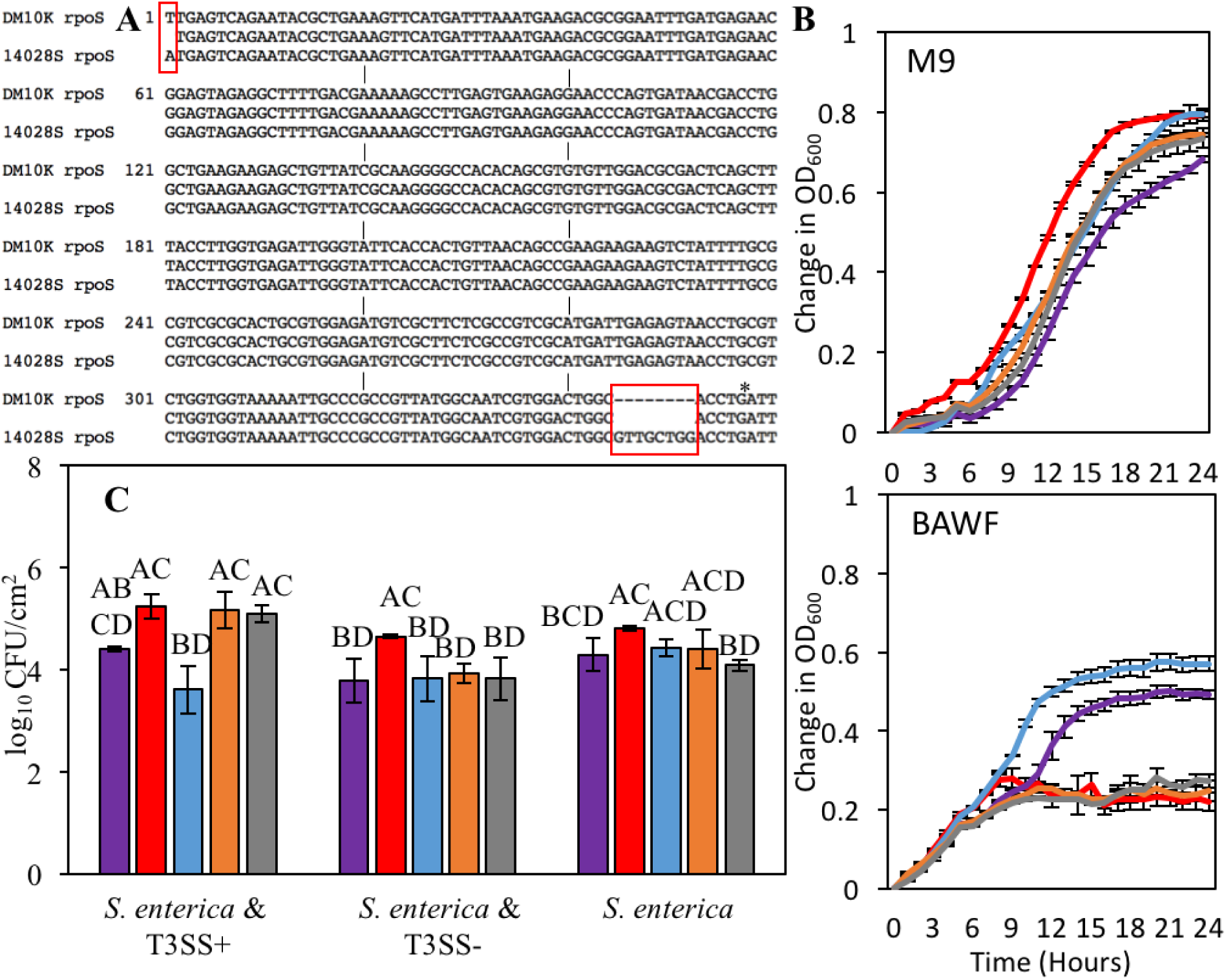
Repairing mutations in the *rpoS* gene of *S. enterica* DM10000 through complementation rescues both *in planta* and *in vitro* phenotypes. **A)** Nucleotide sequence alignment of the first 360 bp of the *rpoS* gene in *S. enterica* strains DM10000 (DM10K) and 14028S. Red boxes indicate nucleotide differences between strains. Asterisk indicates premature stop codon. **B**) Growth curves of 14028S (red), DM10K (purple), DM10KΔrpoS^1-352^ (blue), and DM10KΔrpoS^1-352^::rpoS14028S colonies B1 (orange) and L1 (gray) in M9 minimal media (M9) and filtered *N. benthamiana* apoplastic wash fluid (BAWF). Data was collected and analyzed as described in Fig 3. **C**) Bacterial populations of *S. enterica* strains (same color coding as above) after co-inoculation with *P. syringae* pv. *tomato* DC3000Δ*hopQ1-1* with (T3SS+) and without (T3SS-) a functional Type III Secretion System. Inocula were syringe infiltrated into *N. benthamiana* at concentrations defined in Fig 2. Data are means ± SD (n = 3 plants). Different letters indicate significant differences (2-way ANOVA at p < 0.05).

Despite the complemented strains exhibiting varying colony size, they displayed similar growth to 14028S in both minimal media (M9) and BAWF (Figure 7B). In minimal media, the 14028S, DM10K ΔrpoS^1-352^ deletion, and DM10K complemented strains reached significantly higher final densities compared to the DM10K parental strain. In BAWF, both DM10K and DM10K deletion strains exhibit pronounced biphasic growth with significantly higher densities than the 14028S and DM10K complement strains (Figure 7B). Therefore, the dysfunctional *rpoS* gene in the DM10K strain appears to contribute to the biphasic metabolism of BAWF-derived nutrients, which is compromised when *rpoS* is repaired to a functional gene as in 14028S.

Since *rpoS* restoration in our complement DM10K strains inhibits metabolism of BAWF-derived nutrients, similarly to 14028S, we wanted to determine if our complemented strains behave similarly to 14028S *in planta* during disease. We performed co-inoculation assays for our *S. enterica* parental, deletion, and complement strains in *N. benthamiana* with DC3000 with (T3SS+) a without (T3SS-) a functional T3SS. Both complement strains, DM10KΔrpoS^1-352^::rpoS14028S colonies B1 and L1 displayed significantly greater colonization when co-inoculated with T3SS+ than T3SS- (Fig 7C). Therefore, association with the plant pathogen, DC3K, can promote the growth of our DM10K complement strains. In contrast, 14028S, DM10K, and the *rpoS* fragment deletion strain, DM10KΔrpoS^1-352^, showed no difference in colonization regardless of which strain of DC3000 they were co-inoculated with (Fig 7C). Overall, 14028S has more permissive growth in *N. benthamiana* regardless of its co-inoculation partner compared to DM10K. By itself, the complemented small colony strain, L1, had similar bacterial populations when it was co-inoculated with T3SS-. In contrast, the complemented large colony strain, B1, had similar bacterial populations when it was co-inoculated with T3SS+ (Fig 7C). Overall, our co-inoculation study supports that a functional *rpoS* gene contributes to opportunistic apoplastic colonization by *S. enterica* during plant disease in *N. benthamiana*.

## Discussion

In our experiments, we sought to determine the relevant factors contributing to successful opportunistic endophytic colonization of human enteric pathogens during plant disease. One factor that influences opportunistic growth includes the initial plant pathogen and human enteric pathogen populations. We tested a range of initial inocula with varying O157:H7 and DC3K initial populations to justify our co-inoculation treatments for each host. Generally, a role for the T3SS in the establishment of a more permissive environment for growth of O157:H7 was only observed in *N. benthamiana* with lower concentrations of DC3K, and O157:H7 multiplied to relatively high levels even in the presence of the T3SS-strain. In contrast, a more permissive environment established in *A. thaliana* by the T3SS+ strain compared to the T3SS-strain regardless of initial DC3K populations (Fig 1A). This suggests that there is variability in the general permissiveness of the host apoplast depending on the outcome of host-pathogen interactions, with suppression of PTI playing a more important role in facilitating the growth of enteric pathogens in *A. thaliana* than in *N. benthamiana*. As *N. benthamiana* is not a native host of DC3000, it may elicit a stronger defense response than *A. thaliana*.

We also demonstrated the importance of the ratio of plant pathogen and human enteric pathogen in initial inocula. With 100 fold more O157:H7 than DC3K in the starting inoculum in both model hosts, the growth of DC3K was restricted and a permissive environment could not be established (Fig 1C&D). Hemibiotrophic pathogens such as DC3K share the apoplastic space with human enteric bacteria. This results in potential competition for shared host-derived apoplastic nutrients. We demonstrate that the growth of DC3K is reduced when grown in BAWF with *S. enterica* (Figure 4B). Therefore, an initial inoculum with a significantly greater population of human enteric pathogens than plant pathogens may reduce proliferation of the plant pathogen and thus prevent disease from being established. Another interpretation of this effect is that a greater proportion of human enteric pathogens may overwhelm the plant pathogen population resulting in insurmountable levels of PTI.

In our co-inoculation assays, we found that in most cases when human enteric pathogens such as *E. coli* O157:H7 and *S. enterica* were co-inoculated with the plant pathogenic bacterium, DC3K, with a functional T3SS, the growth of the *E. coli* and *S. enterica* strains was significantly increased compared to populations that were co-inoculated with disarmed DC3K lacking a functional T3SS or *E. coli* and *S. enterica* populations without a co-inoculation partner (Fig 1, Fig S1, Fig 2, Fig 3). This suggests that the *E. coli* and *S. enterica* strains benefit from their association with a plant pathogen partner due to the suppression of plant innate immunity by secreted pathogen effectors. Our results agree with previous studies reporting the beneficial growth of human enteric pathogens from association with bacterial plant pathogens. For example, *S. enterica* exhibits enhanced growth in the phyllosphere of tomato plants during disease caused by the biotrophic pathogen *Xanthamonas perforans* (24). Necrotrophic bacterial plant pathogens that cause soft rots are notorious for enhancing the colonization of both *S. enterica* and *E. coli* in various fresh produce such as lettuce, cilantro, and tomatoes (22, 26, 32).

However, we demonstrate that the permissive environment established by DC3K does not always result in increased growth of human enteric pathogens. In *A. thaliana* and collards, DM10K benefits from co-inoculation with DC3K but not in the host *N. benthamiana* (Fig S1, Fig 2, Fig 3). Contrastingly,14028S benefits from co-inoculation with DC3K in all three hosts (Fig 2C). The ability of 14028S to benefit from DC3K co-inoculation in *N. benthamiana* has been demonstrated previously by Meng *et al.* (23); however, we show that *S. enterica* strain factors are important for opportunistic colonization during plant disease. This suggests that both host and strain factors contribute to the benefit obtained by human enteric pathogens from association with a plant pathogen. With the infectious dose of *Salmonella spp.* and *E. coli spp.* ranging from 10 to 10^5^ bacterial cells, diseased crops could enhance the ability for these human enteric pathogens to reach the infective dose for humans (33, 34, 35).

To elucidate the *S. enterica* strain factors that contribute to differential endophytic colonization during disease in *N. benthamiana*, we sought to determine if DC3K outcompetes DM10K for shared resources in the apoplast. We demonstrated that both *S. enterica* strains are well adapted to metabolize BAWF and M9 minimal media nutrients and DC3K exhibited reduced growth with *S. enterica* strains than by itself in BAWF (Fig 4B). This reduced growth could be due to *S. enterica* strains outcompeting DC3K for shared apoplastic-derived nutrients. Alternatively, the *S. enterica* strains could produce inhibitors during growth in BAWF. Unlike DM10K, 14028S benefitted from co-inoculation with DC3K when grown in BAWF. This suggests that the addition of DC3K to BAWF alters the availability of nutrients that can be utilized by 14028S. Since DM10K did not exhibit the same benefit when co-inoculated with DC3K, this suggests that these two strains differ in their ability to metabolize host-derived nutrients.

We observed differential growth between the two *S. enterica* strains when they were grown in BAWF, but not rich media, minimal media or *A. thaliana* apoplastic wash fluid (Fig 5). More specifically, DM10K exhibited significantly greater growth in BAWF compared to 14028S and both strains exhibited biphasic growth in BAWF. Therefore, we infer that there is a specific compound in BAWF that is differentially metabolized by the two strains. The biphasic growth pattern of the *S. enterica* strains in BAWF, suggest that there are nutrient sources in BAWF that are preferentially metabolized following initial inoculation of bacteria BAWF, and additional nutrient sources that are only metabolized once these preferred nutrients are depleted. To determine whether the observed biphasic growth pattern is linked to catabolite repression, we supplemented BAWF with different macronutrients and micronutrients. As this biphasic growth pattern was suppressed by exogenous glucose and phosphate through catabolite repression, the biphasic growth is most likely due to the metabolism of two or more different carbon sources (Fig 6). However, biphasic growth was either suppressed more in the DM10K strain than the 14028S strain or 14028S repression is alleviated more than DM10K which suggests that these two strains differ in control of catabolite repression of BAWF-derived carbon sources.

Catabolite repression is a tightly regulated process where the presence of a preferred carbon source, such as glucose, inhibits the synthesis of enzymes required to catabolize alternative carbon sources. This process has been extensively studied in *S. enterica* and *E. coli* and is under tight regulation by the cyclic AMP-cAMP receptor protein (CRP) complex (36, 37). Transcriptional regulation of alternative carbon sources is modulated by levels of cAMP which is synthesized by CRP, a global transcriptional regulator. The phosphorylation of EIIA^Glc^ stimulates adenylate cyclase, resulting in the activation of the cAMP-CRP complex which binds to promoter regions for transcription of enzymes required to catabolize alternative sources of carbon. Not only does this complex affect *S. enterica* genes required for growth on numerous carbon sources, but it also affect genes required for virulence, motility, and quorum sensing (38, 39). Exogenous phosphate has previously been shown to suppress catabolite control of the expression of *csr* genes which regulates motility, carbon storage, and virulence in *S. enterica* (39). Our results support a catabolite repression model whereby growth in BAWF is differentially regulated by catabolite repression of BAWF-derived carbon sources in these two *S. enterica* strains.

To determine what types of carbon sources are available in BAWF and which are preferentially catabolized by our strains, we profiled and compared the carbon assimilation abilities of both DM10K and 14028S. We used the phenoarray inhibitor assay to identify which carbon assimilation pathways are constitutively active or induced during growth in BAWF. This technique was established by Rico and Preston (28) who found that DC3K is adapted to use nutrients that are abundant in the tomato apoplast such as trehalose, fructose, galactose, formic acid and citric acid. Based on our un-inhibited phenoarrays data, the metabolic potential for carbon assimilation in our two *S. enterica* strains are very similar with both strains utilizing the majority of the tested carbon sources regardless of what media (LB or BAWF) they were exposed to. In comparing our *S. enterica* un-inhibited phenoarrays results with those published for DC3K, all strains were able to utilize the majority of the carbon nutrients tested of which 60 are shared between the two datasets. However, of these 60 carbon sources, 17 can be utilized by our *S. enterica* strains but not in DC3K. This suggests that these two species may be well adapted for different metabolic niches. For instance, given that our *S. enterica* strains are human enteric pathogens, they are able to utilize mammalian-derived carbon sources such as α-D-lactose and lactulose whereas DC3K, a plant pathogen cannot. Despite our strains being well adapted to the carbon sources available in their specific niches, *S. enterica* strains have the metabolic flexibility to adapt to new niches such as the apoplastic environment.

The inhibitory phenoarray assay was used to identify nutrient assimilation pathways that are constitutively active in a range of medias and pathways that are specifically active in apoplastic-growth *S. enterica* strains. After exposure to a rich media, our inhibited *S. enterica* strains exhibited variations in their ability to metabolize carbon sources with 14028S metabolizing 18 more carbon sources than DM10K despite both strains performing equally well in the uninhibited phenoarrays assays (Table 2). The ability for 14028S to induce the metabolism of more carbon sources may be partially explained by its higher growth rate in LB compared to DM10K (Fig 5A). Since the bacterial subcultures were grown in LB for 3 hours prior to inhibition our cultures may have been at different growth stages and therefore metabolizing the LB-derived carbon sources at different rates. After exposure to BAWF in the inhibition assay, there was variability in the BAWF-induced assimilation pathways between our two *S. enterica* strains with only 4 carbon sources metabolized by DM10K and 49 carbon sources metabolized by 14028S (Table 2). This was unexpected given that, DM0K grows to a higher titer in BAWF compared to 14028S. Given that our subcultures were only grown in BAWF for 3 hours, we can assume that our strains may not have induced all the enzymes required to metabolize BAWF-derived nutrients, especially given that the metabolism of these carbon sources are regulated by catabolite repression. Additionally, we have observed that DM10K has reduced growth during catabolite repression compared to 14028S likely through alleviation of catabolite repression in 14028S or stronger repression in DM10K (Fig 6). Thus we hypothesize that this regulatory process may explain the lack of BAWF-induced carbon metabolism observed in DM10K. Performing our inhibition phenoarrays assay at a later time point during growth in BAWF may broaden the availability of BAWF-induced carbon metabolizing enzymes.

Apoplastic-grown DM10K used two metabolites that were not used by 14028S, 1,2 propanediol and glycolic acid. The organic compound 1,2 propanediol or propylene glycol is a part of the proponoate metabolism pathway and is utilized in pharmaceuticals according to its KEGG compound profile (C00583). Glycolic acid or glycolate is a byproduct of the photorespiration system in plants and can be utilized to form the amino acids serine and glycine (40). Although none of these products were detected by GS-MS in BAWF, the enzymes required to metabolize similar BAWF-derived carbon structures may have been activated in DM10K likely though the glycerate or glyoxylate shunt pathways. Apoplastic-grown 14028S, after inhibition treatment, utilized 52 carbon sources, and therefore gives a more comprehensive insight as to what carbon sources may be available in BAWF. Of these 52 carbon sources, 14028S induced the metabolism of 11 sugar or sugar-derivatives including trehalose, glucose, fructose, ribose, galactose, mannose and cellobiose (Table 2). Trehalose, fructose, galactose, mannose, and glucose have been previously identified to be metabolized by *Pseudomonads* using similar methods in both tomato and bean apoplastic wash fluid (28, 41). Cellobiose is the disaccharide form of cellulose and is classified as a plant metabolite according to its ChEBI profile (CHEBI:17057). The *S. enterica* genome encodes multiple phosphotransferase systems (PTS) to transport and phosphorylate a number of sugar substrates including glucose, mannose, fructose, trehalose and cellobiose (*cel* operon). (42). Glucose, mannose, and trehalose have been found to be regulated by CRP-cAMP for catabolite repression in bacteria and therefore some sugars may play a role in the observed biphasic growth in BAWF (42, 43). GC-MS analysis of AWF from bean during infection by *P. syringae* pv. *phaseolicola* demonstrate that the bacteria preferentially metabolizes malate, glucose, and glutamate while excluding abundant apoplastic metabolites such as citrate and GABA until the preferred metabolites were depleted (41). Therefore, this catabolite repression of AWF-derived nutrients have been previously demonstrated by bacteria occupying the plant apoplast.

Our GC-MS analysis of BAWF was able to identify various sugars, amino acids and sugar alcohol derivatives (Table S2). Of the 30 shared compounds between the PM1 BIOLOG plate and our GC-MS data, 20 compounds were identified to be metabolized by 14028S which suggests that our GC-MS analysis supports our findings from our inhibitory phenoarrays assay. Of these 20 compounds, 11 have been previously identified to be metabolized by bacteria grown in tomato apoplastic wash fluid include succinic acid, L-aspartic acid, L-glutamic acid, L-serine, L-asparagine, L-alanine, D-fructose, D-glucose, D-galactose, D-trehalose, and glutaric acid (28). Due to the limited carbon sources in our phenoarray assay, repeating this assay with additional carbon sources will give us a more comprehensive understanding of what carbon sources are available in BAWF.

Given that our *S. enterica* strain DM10K is an LT2 derivative with a known start codon mutation in its *rpoS* gene, we sought to determine if this mutation also contributes to the phenotypic differences we observed between our DM10K and 14028S strains. Upon sequencing the *rpoS* gene in our DM10K strain, we observed the alternative TTG start codon as well as an additional 8 bp deletion resulting in a premature stop codon (Fig 7A). Therefore, DM10K most likely has a truncated RpoS protein of 117 aa compared to 14028S whose *rpoS* gene encodes a fully functional 330 aa protein. RpoS (σ38, σ^S^) is a sigma factor with a known regulon. RpoS is the master regulator of the general stress response, which is triggered by many different stress signals resulting in either a reduction of growth or aids in the survival and protection against additional stressors (44). Low levels of RpoS, as a result from the alternative start codon in LT2, contributes to the strains avirulent phenotype in mice by altering the expression of virulence genes found in the RpoS regulon (31). To determine whether the mutations in the DM10K *rpoS* gene contribute to the observed growth phenotypes both in BAWF and *in planta*, we deleted the first 352 bp in *rpoS* containing the mutations. We complemented this DM10KΔrpoS^1-352^ mutant via allelic exchange with the first 352 bp of *rpoS* from 14028S to restore the full length *rpoS* gene in our DM10KΔrpoS^1-352^::rpoS14028S complement strain. Our complement strain had various colony morphologies with small colonies similar in size to DM10K and large colonies similar in size to 14028S (Fig S6). Analysis of the RpoS regulon demonstrates that RpoS is directly involved in the expression of genes involved in the biogenesis and structure of the LPS and outer membrane proteins (45). It is possible that variations in the *rpoS* gene contribute to the observed differences in colony size between our strains. However, our DM10KΔrpoS^1-352^ mutant strain had significantly larger colony diameter than its parental strain DM10K and similar colony size to the fully functional *rpoS* strain 14028S (Fig S6). This suggests that the variation in the *rpoS* gene does not play a role in colony size. Variations in the complement strain colony size, despite having confirmed sequences for the *rpoS* gene and promoter region, may be explained by differential regulation at the translational or protein level.

Despite the variation in colony size in our complement strain, both colony morphologies exhibit similar phenotypic growth to 14028S in BAWF (Fig 7B). More specifically, the RpoS restored strains exhibit reduced biphasic growth compared to the DM10K and DM10KΔrpoS^1-352^ mutant strains. Additionally, our DM10KΔrpoS^1-352^ mutant strain exhibited greater overall growth in BAWF compared to its parental DM10K strain which suggests that the truncated RpoS protein in DM10K has reduced, but possibly not abolished, gene regulation. Our results support the role of RpoS in regulating the metabolism of BAWF-derived carbon sources in our two *S. enterica* strains. It has been previously demonstrated that mutations in *rpoS* accumulate during stationary phase and in glucose-limiting conditions in *E. coli* in order to improve nutrient scavenging under nutrient starved conditions (30, 46, 47). Therefore, DM10K is better adapted to metabolize BAWF-derived sugars, as low levels of RpoS allow it to metabolize a more diverse range of nutrient sources especially under nutrient starved conditions. Our BIOLOG analysis did not support this hypothesis in that the number of BAWF-induced carbon substrates metabolized by 14028S following inhibitor treatment was far greater than that of DM10K. One explanation for this may be due to the catabolite repression response in DM10K during the early stages of growth in BAWF.

King *et al*. utilized the un-inhibited carbon phenoarrays on *E. coli* strains with low and high levels of RpoS and showed that *rpoS* disruption allows for the stimulation of more carbon substrates including D-melibiose, B-methyl-D-glucoside, L-rhamnose, D-sorbitol, acetic acid, D-galacturonic acid, succinic acid, bromosuccinic acid, L-alanine, L-alanyl-glycine, L-asparagine, L-aspartic acid, and DL-glycerol phosphate (30). The majority of these compounds were metabolized by both DM10K and 14028S grown in both LB and BAWF in our un-inhibitory phenoarrays assay. Additionally, the majority of these compounds exhibited BAWF-induced metabolism by 14028S and therefore could be potential BAWF-derived carbon sources that are differentially metabolized by our two *S. enterica* strains. Transcriptome and phenoarray analysis of *E. coli rpoS* and *cya* mutants reveals that the absence of cAMP and not RpoS has a negative impact on the transcription of catabolic genes for alternative carbon substrates during growth in glucose-limiting media. More specifically, *rpoS* mutants exhibited reduced rates of oxidation of trehalose, mannitol, sorbitol, and D-malate in inhibited phenoarrays assays in glucose-limited media (48). In our inhibited phenoarrays analysis, there were no differences between the ability of DM10K and 14028S to metabolize trehalose and mannitol after 3 hours of exposure to LB (Table 2). If there are still high levels of glucose in BAWF by 3 hours when samples were taken, low levels of cAMP may contribute to the lack of metabolism of carbon sources especially if catabolite repression is stronger in our DM10K strain than 14028S. Expansion of our phenoarrays analysis during the various phases of growth in BAWF would need to be performed in order to address this hypothesis. Although we demonstrate that *rpoS* plays a role in regulating metabolism of BAWF-derived carbon sources, cAMP-CRP is a regulator of *rpoS* transcription depending on the bacterial growth phase with two putative cAMP-CRP binding sites in the promoter region of *rpoS* in *E. coli* (44).

Not only did our complement strains exhibit reduced growth in BAWF, they exhibited improved colonization during disease in *N. benthamiana* similarly to 14028S (Fig 2C, 7C). In contrast, both DM10K and the DM10K *rpoS* mutant strains did not benefit from co-inoculation with T3SS+. This suggests that *rpoS* plays a role in opportunistic *in planta* colonization during plant disease. The plant innate immune response causes various changes in the apoplast that make it less habitable to invading bacteria through the production of reactive oxygen species (ROS) and phytoalexins, diversion of water away from the apoplast, changes in pH, and membrane polarization (17, 48, 49). DC3K creates a more favorable apoplastic environment for colonization by suppressing these changes through the delivery of effectors (17, 20, 50, 51). Environmental stressors including high osmolarity, heat shock, acidic pH, and oxidation induce elevated intracellular RpoS levels as *rpoS* plays a crucial role in stress tolerance in bacteria (52, 53, 54, 55). With the documented mutations in *rpoS* in our DM10K strain, we attribute its inability to colonize the apoplast of *N. benthamiana* during infection by DC3K to its reduced tolerance to plant-derived stressors including reduced pH, high osmolarity and ROS production in the apoplast. Similar results were observed in *Medicago truncatula* where 14028S was more tolerant than LT2 to the FLS2-dependent immune response which was attributed to its RpoS-dependent response to oxidative stress (55). In our system, DC3K suppresses the immune response by *N. benthamiana* which in turn should reduce the different plant-derived stressors; however, the suppression may only be sufficient for DC3K colonization who is likely to be more tolerant than *S. enterica* strains due to its co-evolution with plants.

In summary, our study identifies both plant and strain factors that contribute to the opportunistic apoplastic colonization of human enteric pathogens during plant disease by conducting co-inoculation assays in various plant hosts with different human enteric pathogens strains and the plant pathogen DC3K. Variations in plant defense response and nutrient availability in the apoplast between hosts are likely some of the host factors important for colonization by human enterics. Our human enteric strains exhibited variable growth in apoplastic wash fluid collected from different plant hosts which suggest that some strains are better suited to metabolize plant-derived nutrients within the apoplast. We demonstrate that RpoS plays an important role in regulating the metabolism of plant-derived nutrients as strains with low levels of RpoS have been found to be more competitive in carbon-limiting environments. However, there is a fitness cost to this expanded nutritional capacity in that strains with low levels of RpoS are likely to have reduced resistance to apoplastic stressors such as osmotic stress, oxidative stress, and low pH. This fitness trade-off has been previously documented in *E. coli* where nutrient limitation was found to increase selection pressure for loss of *rpoS* functionality, but low pH and high osmolarity reduced fitness in strains with reduced rates of *rpoS* enrichment (56). As RpoS levels play an important role in the survival of human enteric pathogens within the environment, monitoring polymorphisms in *rpoS* within plant-associated populations could provide some fruitful information on how to mitigate produce-borne outbreaks of human enteric pathogens.

## Methods

### Plant Tissue and Bacterial Culture Preparation

*A. thaliana* Col-0 seeds suspended in sterile 0.1% agarose were sown in SunGrow Professional potting mix and stratified in darkness for 1 day at 4°C before being grown in a growth chamber (Conviron A1000) with 14-h light (70 μmol) at 23°C. Plants were removed from the chamber at 4 weeks and kept at 12-h day and 12-h night conditions in the growth room prior to inoculation (4-5 weeks old) or apoplastic extractions (6 weeks or older). *N. benthamiana* and collard (*B. oleracea* var. *acephala* cv. Morris Heading) plants were sown in the same potting mix amended with 1g/L Peter’s 20-20-20 fertilizer and grown in a growth chamber with 12h day at 26°C (70 μmol) and 12h night at 23°C. Two weeks after sowing, seedlings were transplanted into 6 inch pots and fertilized. Plants were removed from the chamber at 5 weeks and kept in the growth room prior to inoculations or apoplastic extractions (6-9 weeks old). Collard seedling inoculations were conducted 3 weeks after sowing. For metabolomics analyses *Nicotiana benthamiana* plants were grown at 22 °C and 60 % relative humidity under a 12 h light regime. Leaves from 4-week-old N. benthamiana plants were used for apoplastic fluid isolation.

*P. syringae* pv. *tomato* strain DC3000 (DC3K), isogenic mutant derivations, *S. enterica* serovar Typhimurium strains, and *E. coli* O157:H7 strain used in this study are listed in Table 1. All DC3K strains were grown on King’s B medium with 60 μg mL^-1^ of rifampicin at 30°C. All *E. coli* and *S. enterica* strains were grown on Luria-Bertani (LB) medium with 50 μg mL^-1^ of kanamycin (*E. coli* only) at 37°C. To enumerate *S. enterica* populations from a mixed population, samples were grown at 42°C.

### Plant Inoculation and Sampling Procedure

To prepare *E. coli* and *S. enterica* bacterial inocula, overnight cultures made from single colonies were incubated in LB at 37°C were pelleted using centrifugation, suspended in 0.25 mM MgCl_2_, and were diluted to an optical density at 600 nm (OD_600_) of 0.8 (approximately 5 x 10^8^ CFU/mL), as determined using a Biospectrometer (Eppendorf, Hamburg, Germany). DC3K inoculum was prepared as described in Lovelace *et al*. (57) and diluted to OD_600_ = 0.8. Bacterial inocula for both individual and co-inoculations were further diluted in 0.25 mM MgCl_2_ to the desired concentrations. *E. coli* at 5 x 10^4^ CFU mL^-1^ and *S. enterica* strains at 5 x 10^5^ CFU mL^-1^, were mixed with DC3K (pathogenic to *A. thaliana*) or DC3KΔ*hrcC* (a type III secretion mutant) to a final DC3K concentration of 5 x 10^6^ CFU mL^-1^ and syringe-inoculated into four *A. thaliana* leaves per plant. *E. coli* at 5 x 10^6^ CFU mL^-1^ and *S. enterica* strains at 5 x 10^5^ CFU mL^-1^, were mixed with DC3KΔ*hopQ1-1* (compatible with *N. benthamiana*) or DC3KΔ*hrcC* to a final DC3K concentration of 5 x 10^4^ CFU mL^-1^ or 5 x 10^5^ CFU mL^-1^ for *E. coli* and *S. enterica* co-inoculations respectively and syringe inoculated into fully expanded *N. benthamiana* leaves. *E. coli* and *S. enterica* strains were mixed with DC3K (pathogenic to collards) or DC3KΔ*hrcC (not pathogenic)* to a final concentration of 5 x 10^5^ CFU mL^-1^ and syringe-inoculated into fully expanded collard leaves. These inocula were used for single or co-inoculations in all plant hosts unless otherwise noted. All inoculated hosts were incubated for three days under high humidity (90%-100% RH) unless otherwise noted.

For spray inoculations on collard seedlings, DC3K strains were diluted to an OD_600_ of 0.8 (5 X10^8^ CFU mL^-1^) in 0.02% Silwet solution, sprayed onto seedlings, and incubated for five days under high humidity. Infiltrated leaves were sampled at different time points, homogenized in 0.25 mM MgCl_2_, and serial dilutions were plated on appropriate plates with antibiotics to determine the population size of both enteric strains and DC3K strains measured as CFU cm^-2^. All plant inoculation assays were repeated three times with 3-4 plants per treatment.

### Extraction of Apoplastic Wash Fluid, BIOLOG phenoarrays, and GC-MS

Apoplastic wash fluid (AWF) was crude extracted using vacuum infiltration as described by O’Leary *et al*. (58) with slight modifications. Whole *A. thaliana* plants or fully expanded *N. benthamiana* leaves were cut and placed into a 500 mL beaker with 300 mL of distilled water. Repeated cycles of vacuum at 95 kPa for 2 min followed by slow release of pressure were applied until leaves were fully infiltrated. Excess water was blotted from plant tissue before leaves were rolled into 20 mL syringes which were placed into 50 mL conical tubes. Tubes were centrifuged at 1,000 rpm for 10 min at 4°C and the fractions were pooled and stored at −80°C. AWF samples were filter sterilized using 0.2 μM RapidFlow filters for subsequent experiments. The crude extractions were not measured for cytoplasmic contamination.

For comparative analysis of *S. enterica* strains’ ability to use a range or compounds as carbon sources after pre-treatment in *N. benthamiana* apoplastic wash fluid (BAWF) or LB, Biolog PM1 plates were inoculated using methods defined by Rico and Preston (28) and following the manufacturer’s instructions (BIOLOG, Hayward, CA, U.S.A) with modifications. Bacterial inocula were generated by overnight culture in LB for DM10K and 14028S. A 1 mL aliquot of each strain was washed twice with 0.25mM MgCl_2_ and resuspended in 5 mL of BAWF or LB. Samples were incubated with shaking at 37°C for 3 hours. To remove excess carbon source, samples were centrifuged and washed twice with 0.25 mM MgCl_2._ Washed cells were resuspended in 10 mL of 1X IF-0a GN/GP Base inoculating fluid containing 1X Redox Dye Mix A (BIOLOG, Hayward, CA, U.S.A) to a final OD_600_ of 0.3 ± 0.05. Aliquots of 100 μL were inoculated into each well of the Biolog PM1 plate. Each plate was read using a Tecan Spectra Rainbow microplate reader (Tecan, Männedorf, Switzerland) and initial absorbance values for OD_460_ was recorded for all wells. Absorbance values were normalized by subtracting from the negative control value. Plates were incubated with shaking at 37°C for 24 hours. Final absorbance values for OD_460_ was recorded for all wells and normalized to the negative control well. The change in OD_460_ was calculated for each well by subtracting the initial normalized OD_460_ from the final normalized OD_460_.

To inhibit expression of new proteins during incubation in the PM1 plates, bacterial samples were treated with 10 μg/μL tetracycline after 3 hours of incubation in either LB or BAWF. After inhibition treatment, samples were centrifuged and washed twice with 0.25 mM MgCl_2_ containing 10 μg/μL tetracycline to remove excess carbon sources. Washed cells were resuspended in 10 mL of 1X IF-0a GN/GP Base inoculating fluid containing 1X Redox Dye Mix A and 10 μg/μL tetracycline (BIOLOG, Hayward, CA, U.S.A) to a final OD_600_ of 0.3 ± 0.05. The inhibited samples were inoculated onto PM1 plates and incubated as described above in the un-inhibited samples. Phenoarrays were repeated twice and average change in normalized absorbance values were evaluated.

BAWF samples were collected as described by O’Leary *et al*. (41). BAWF samples were prepared for GC-MS analysis using a modified version of the method of Lisec *et al.* (59). One hundred and fifty microliter samples of BAWF were mixed with 700 μL of methanol, supplemented with 10μg/mL of ribitol, and then shaken at 70°C for 10 minutes, followed by centrifugation for 5 minutes at 11000 g. Next, 700 μL of supernatant was removed and mixed sequentially by vortexing with 375 μL of cold chloroform and 500 μL of cold ddH_2_O. Samples were centrifuged at 2200 g for 15 minutes, then 250 μL of supernatant was transferred to a fresh tube and dried in a vacuum concentrator without heat. The samples were derivitized in 29 μL of pyridine containing 20 mg/mL methoxyamine and 50 μL of N-Methyl-N-(trimethylsilyl) trifluoroaceamide (MSTFA) as described previously (59). GC-MS analysis was performed as described previously (41).

### Bacterial Growth Assays

Bacterial inocula were generated as described above for all strains. Cell suspensions in 0.25 mM MgCl_2_ were standardized to an OD_600_ of 0.8 (5 x 10^8^ CFU mL^-1^). Aliquots of 300 μL of bacterial inocula were diluted into 2.7 mL of BAWF, LB, or M9 minimal media (42) to a final concentration of approximately 5 x 10^7^ CFU mL^-1^. To make macronutrient and micronutrient amended BAWF, 1000X concentrated macronutrients and micronutrients including sodium chloride, magnesium sulfate, ammonium sulfate, calcium chloride, glucose, potassium phosphate, potassium chloride, and sodium phosphate, were dissolved in distilled water and filter sterilized using 0.2 μM filters before being diluted to 1X in BAWF. The same volume of water was used as a control. Aliquots of 400 μL were inoculated into 5 replicate wells of a Bioscreen honeycomb plate (Bioscreen Technologies, Bertinoro, Italy). The OD_600_ was measured every hour for up to 36 hours in a Bioscreen C plate reader with low to medium shaking at 22°C (Bioscreen Technologies, Bertinoro, Italy). Raw absorbance readings were normalized by subtracting the initial absorbance readings from subsequent hourly readings.

For the competition assay, standardized inocula of each *S. enterica* strain and DC3K were diluted in combination with either 0.25 mM MgCl_2_ or an inoculum partner for individual or co-inoculations respectively in M9 minimal media or apoplastic wash fluid to a final concentration of 5 x 10^6^ CFU mL^-1^ of each bacterium. Aliquots of 400 μL were inoculated into 5 replicate wells of a Bioscreen honeycomb plate and the growth (measured as OD_600_) was monitored until peak OD_600_ was achieved; one day for samples grown in BAWF and two days for samples grown in minimal media. Serial dilutions were plated on appropriate plates with antibiotics to determine the initial and final population sizes of both *S. enterica* strains and DC3K measured as CFU mL^-1^. Initial populations were measured from three aliquots of the original suspension and final populations were measured from individual wells in the Bioscreen honeycomb plate. At least two independent experiments were performed for all growth assays.

### Gene Fragment Swap by Allelic Exchange

*S. enterica* knock-out clones were generated in the DM10K strain background using the pR6KT2G suicide vector which allows for SacB-mediated sucrose counter-selection using methods defined by Stice *et al*. (60) with modifications. The promoter and gene sequence of *rpoS* (STM14_3526) from 14028S was obtained from KEGG Gene using the organism code “seo”. The *rpoS* gene fragment to be deleted is the first 352 bp of the gene. Flanks of 300 bp preceding and following the gene fragment were synthesized with *attB1* and *attB2* extensions for Gateway compatibility as double stranded DNA gblocks by Twist Bioscience (Table S1). The synthesized gene fragment was Gateway cloned into pR6KT2G through a BP clonase reaction according to the manufacturer’s protocol. (Thermo Scientific, Waltham, MA, USA). The cleaned reaction mixture was electroporated into competent *E. coli* MaH1 *pir*^+^ cells and transformed cells were grown on LB amended with 10 μg/μL gentamycin. Knock-out *rpoS* constructs were confirmed by BsrGI digest and sequencing before transformed into electrocompetent *E. coli* RHO5 *pir*^+^ cells.

The wild-type DM10K strain and RHO5 pR6KT2G:*rpoS* donor strain were mated on LB plates amended with 250 μg/μL Diaminopimelic Acid (DAP). Merodiploids were recovered from the mating mixture on LB plates amended with 10 μg/μL gentamycin. Two merodiploid colonies were selected for counter selection in a liquid culture of 1mL LB and 3mL 1M sucrose for 24 hours at 37°C. Following counter selection, a portion of the diluted mixture was plated on LB plates amended with X-gluc. Candidate knock-out strains were not blue in color and thus evicted the plasmid construct. Genomic DNA extractions were performed on candidate colonies using the Gentra Puregene kit according to the manufacturer’s instructions (Qiagen, Hilden, Germany). Candidate colonies were screened using external primers (Table S1) by PCR using Phusion HiFi polymerase according to the manufacturer’s instructions (New England BioLabs, Ipswich, MA, USA). Candidates with the expected PCR fragment size were sequenced using external primers to confirm the knock-out of the gene fragment.

The resulting DM10KΔrpoS^1-352^ strain was complemented with the *rpoS* 352bp gene fragment from 14028S using the same homologous recombination procedure used to generate the mutants. Flanks of 300 bp preceding and following the 14028S *rpoS* gene fragment were synthesized with *attB1* and *attB2* extensions for Gateway compatibility as double stranded DNA gblocks by Twist Bioscience (Table S1). This gene fragment was cloned as described above however, transformed cells and resulting matings were plated on LB amended with 10 μg/μL gentamycin. Candidate knock-in strains were screened using external primers by PCR; the resulting gene fragment was digested using the AvaII restriction enzyme according to the manufacturer’s instructions to distinguish 14028S and DM10K genotypes (New England BioLabs, Ipswich, MA, USA). Candidates with the expected PCR and digest fragment sizes were sequenced using the out primer and promoter primer to confirm the 14028S *rpoS* gene knock-in clones. Confirmed complement strains, deletion strain, and parental strains DM10K and 14028S were streaked to isolation from an overnight culture in LB and incubated at 37°C overnight. Colonies from each plate were imaged using a Nikon camera with a ruler to scale. Images were loaded into ImageJ v.2.1.0 and the scale was set using the line segment tool to span 2 mm on the photographed ruler. Using the line segment tool, five colony diameters were measured from each plate and recorded.

## Acknowledgements

The authors thank Jinru Chen and Anna Glasgow Karls (University of Georgia) for providing strains, Samantha Ayoub (University of Georgia) for performing spray inoculations on collards and the members of Brian Kvitko’s and Li Yang’s (University of Georgia) labs for assistance in reviewing the manuscript.

---

**Fig S1.**
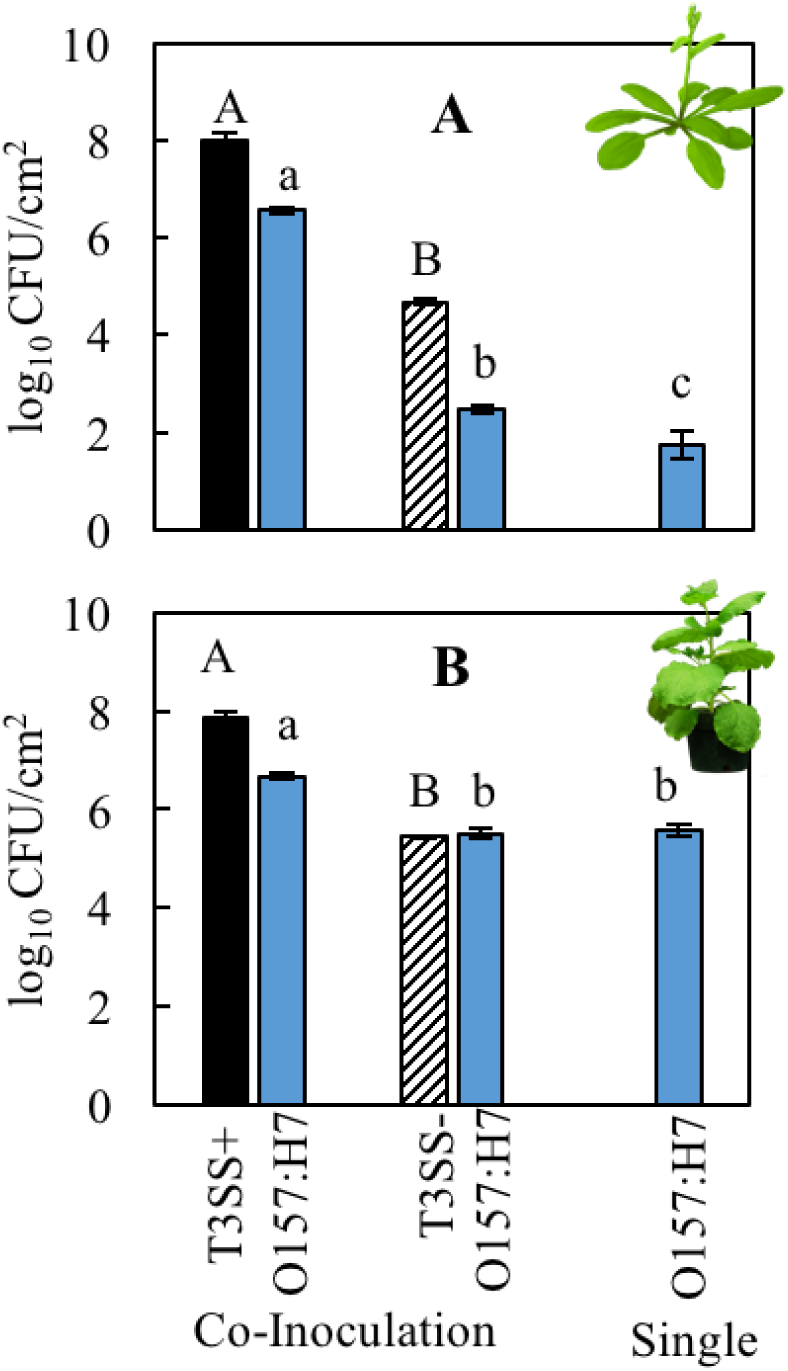
Increased endophytic colonization of *E. coli* O157:H7 during plant disease established by *P. syringae* in two model hosts. Bacterial populations of *E. coli* O157:H7 5-11 (O157:H7, blue) with co-inoculation partners *P. syringae* pv. *tomato* DC3000 or DC3000Δ*hopQ1-1* with (T3SS+, black) and without (T3SS-, striped) a functional Type III Secretion System. Inocula were syringe infiltrated into model plant hosts, **A**) *A. thaliana* Col-0 at a concentration of 5 x 10^6^ CFU mL-1 for DC3000 strains and 5 x 10^4^ CFU mL^-1^ *E. coli* and **B**) *N. benthamiana* at a concentration of 5 x 10^4^ CFU mL-1 for 1122 DC3000 strains and 5 x 10^6^ CFU mL^-1^ *E. coli*. Bacterial populations were measured as log colony forming units per cm2 of leaf tissue (log^10^ CFU/cm^2^) 3 days post-inoculation. Data are means ± SD (n = 3 plants). Different letters indicate significant differences (2-tailed t-test for each strain at p < 0.05).

**Fig S2.**
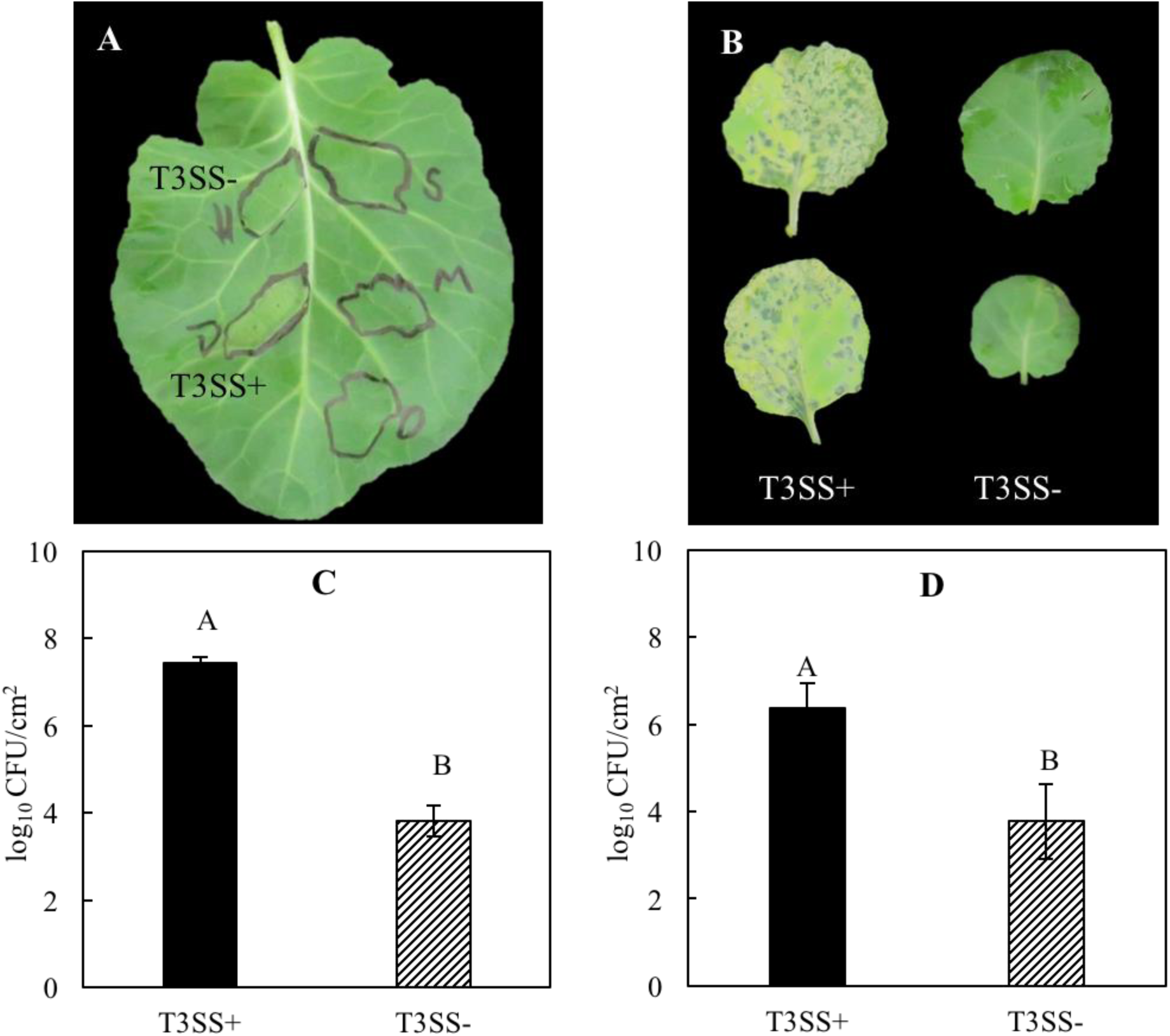
*P. syringae* can cause disease on collard leaves in a T3SS dependent manner. Disease symptoms of *Brassica oleracea* var. *acephala* (collards) after infection with *P. syringae* pv. *tomato* DC3000 with (T3SS+) and without (T3SS-) a functional Type III Secretion System after **A**) syringe and **B**) spray inoculation. T3SS+ and T3SS- were syringe inoculated into adult collard leaves at a concentration of 5 x 10^5^ CFU mL^-1^ or spray inoculated onto seedlings at a concentration of 5 x 10^8^ CFU mL^-1^. Disease symptoms were observed 3 and 5 days post inoculation (dpi) respectively. **C**) Bacterial populations in adult collard leaves 3 dpi and **D**) in collard seedlings 5 dpi. Bacterial populations were measured as log colony forming units per cm^2^ of leaf tissue (log_10_ CFU/cm^2^). Data are means ± SD (n = 3 plants). Different letters indicate significant differences (2-tailed t-test at p < 0.05).

**Fig S3.**
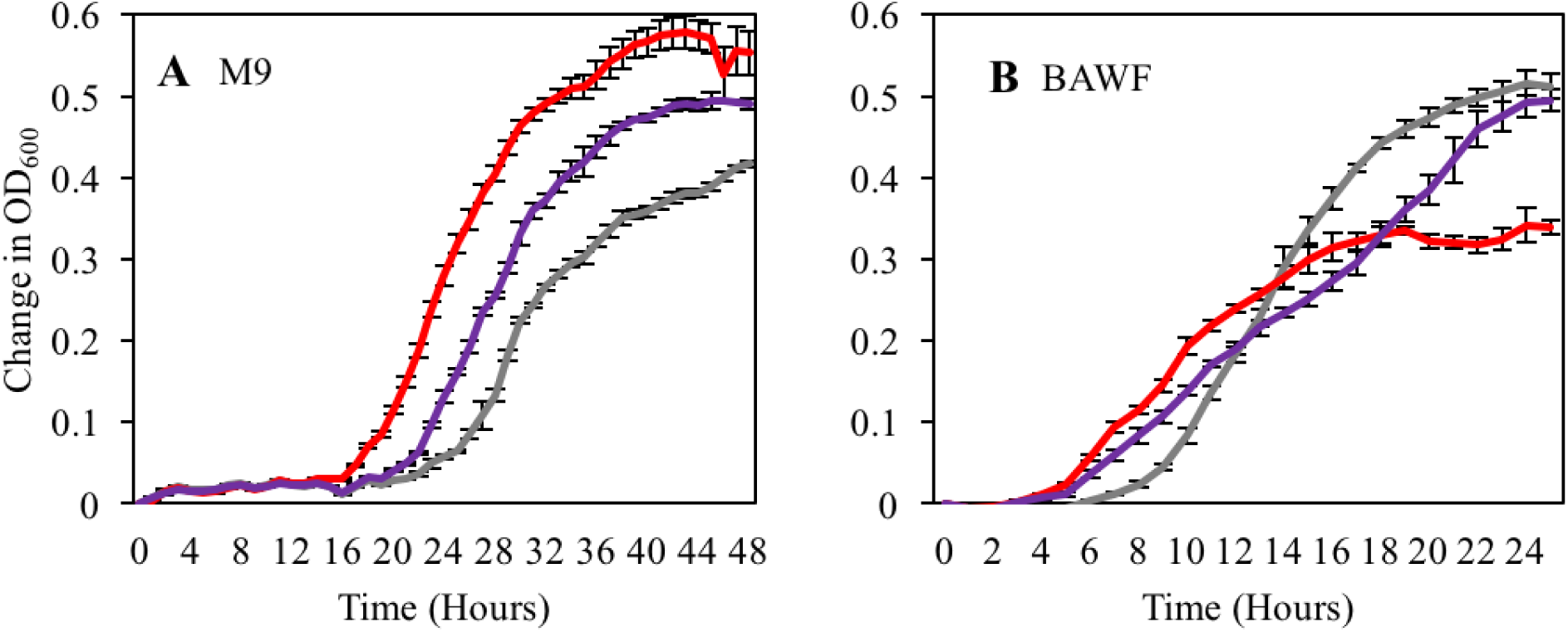
Growth of *S. enterica* and *P. syringae* strains in minimal media and *N. benthamiana* apoplastic wash fluid. Growth curves of *S. enterica* DM10000 (purple), *S. enterica* 14028S (red), and *P. syringae* pv. *tomato* DC3000 (gray) in **A**) M9 minimal media and **B**) filtered *Nicotiana benthamiana* apoplastic wash fluid. Cultures were aliquoted into 5 replicate wells, incubated at 22°C, and the OD_600_ was recorded every hour. Growth was measured as the average change in measured OD_600_ with standard deviation error bars (n=5 wells).

**Fig S4.**
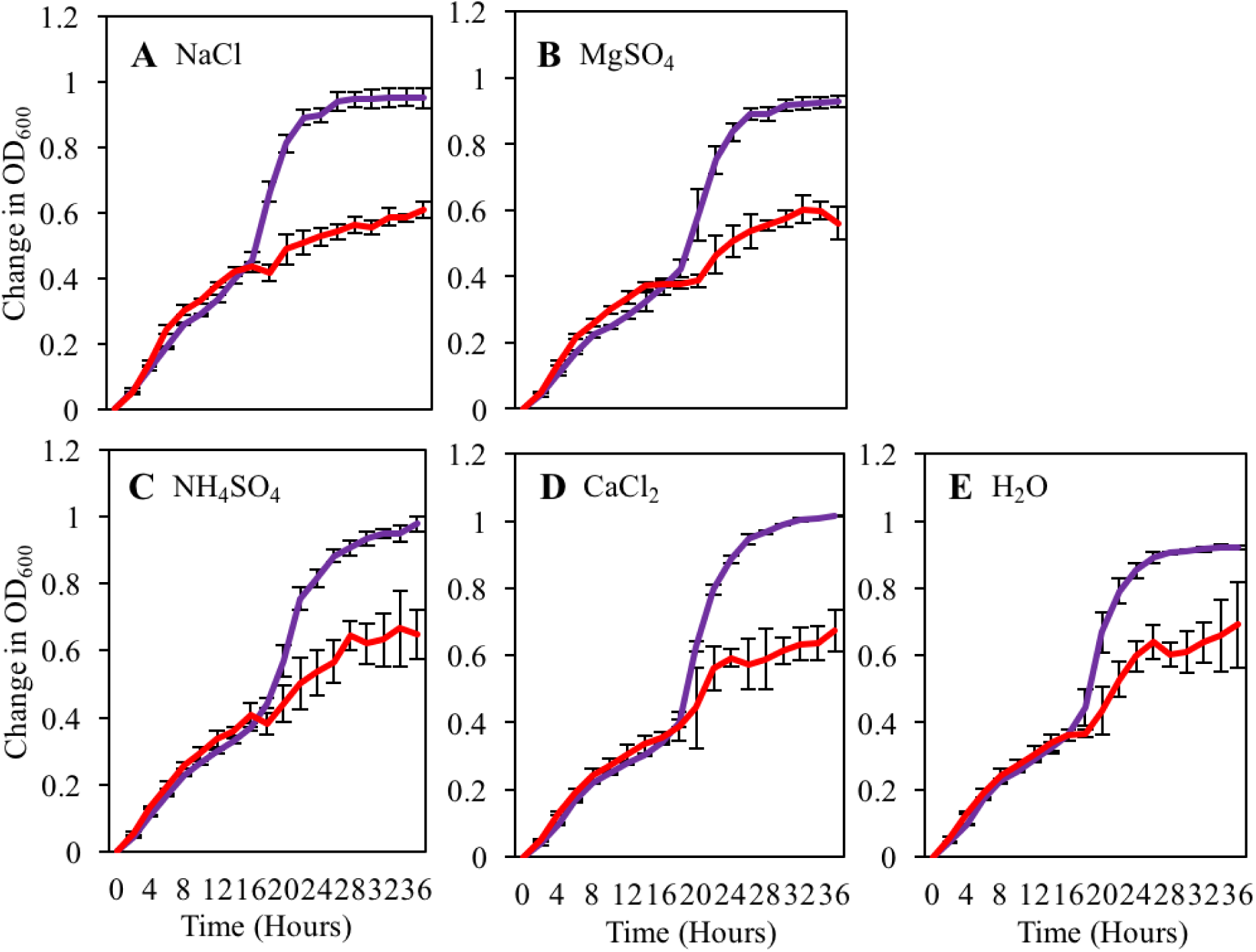
*S. enterica* biphasic growth in *Nicotiana benthamiana* apoplastic wash fluid is unaltered after supplementation with specific macro and micronutrients. Growth curves of *S. enterica* DM10000 (purple) and *S. enterica* 14028S (red) in filtered *N. benthamiana* apoplastic wash fluid supplemented with **A**) 10 mM sodium chloride, **B**) 5 mM magnesium sulfate, **C**) 10 mM ammonium sulfate, **D**) 1 mM calcium chloride, and **E**) water. Cultures were aliquoted into 5 replicate wells, incubated at 22°C, and the OD_600_ was recorded every 2 hours. Growth was measured as the average change in measured OD_600_ with standard deviation error bars (n=5 wells).

**Fig S5.**
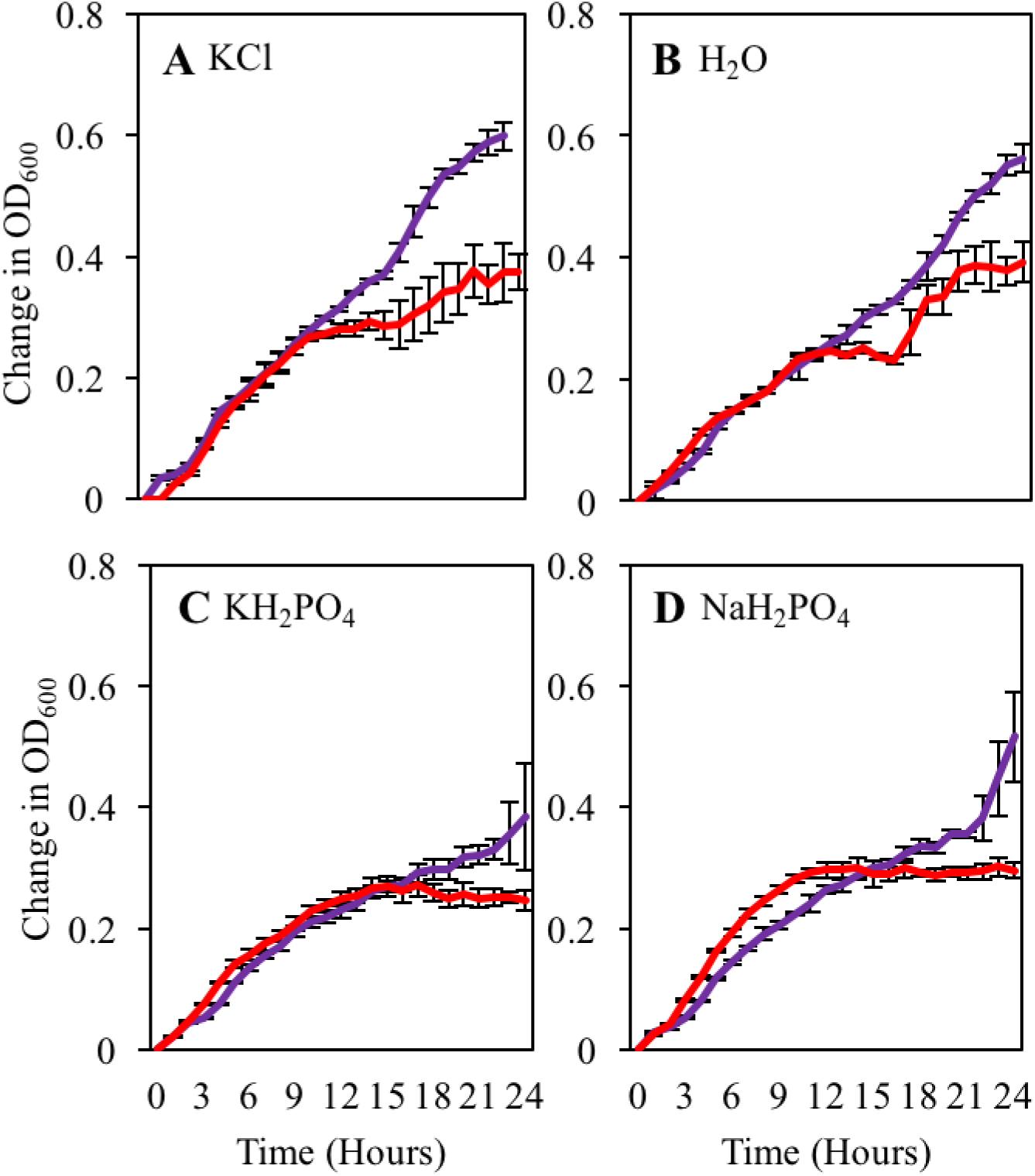
*S. enterica* biphasic growth in *Nicotiana benthamiana* apoplastic wash fluid is suppressed by the phosphate anion. Growth curves of *S. enterica* DM10000 (purple) and *S. enterica* 14028S (red) in filtered *Nicotiana benthamiana* apoplastic wash fluid supplemented with **A**) 10 mM potassium chloride, **B** water, **C**) 10 mM potassium phosphate, and **D**) 10 mM sodium phosphate. Cultures were aliquoted into 5 replicate wells, incubated at 22°C, and the OD_600_ was recorded every hour. Growth was measured as the average change in measured OD_600_ with standard deviation error bars (n=5 wells).

**Fig S6.**
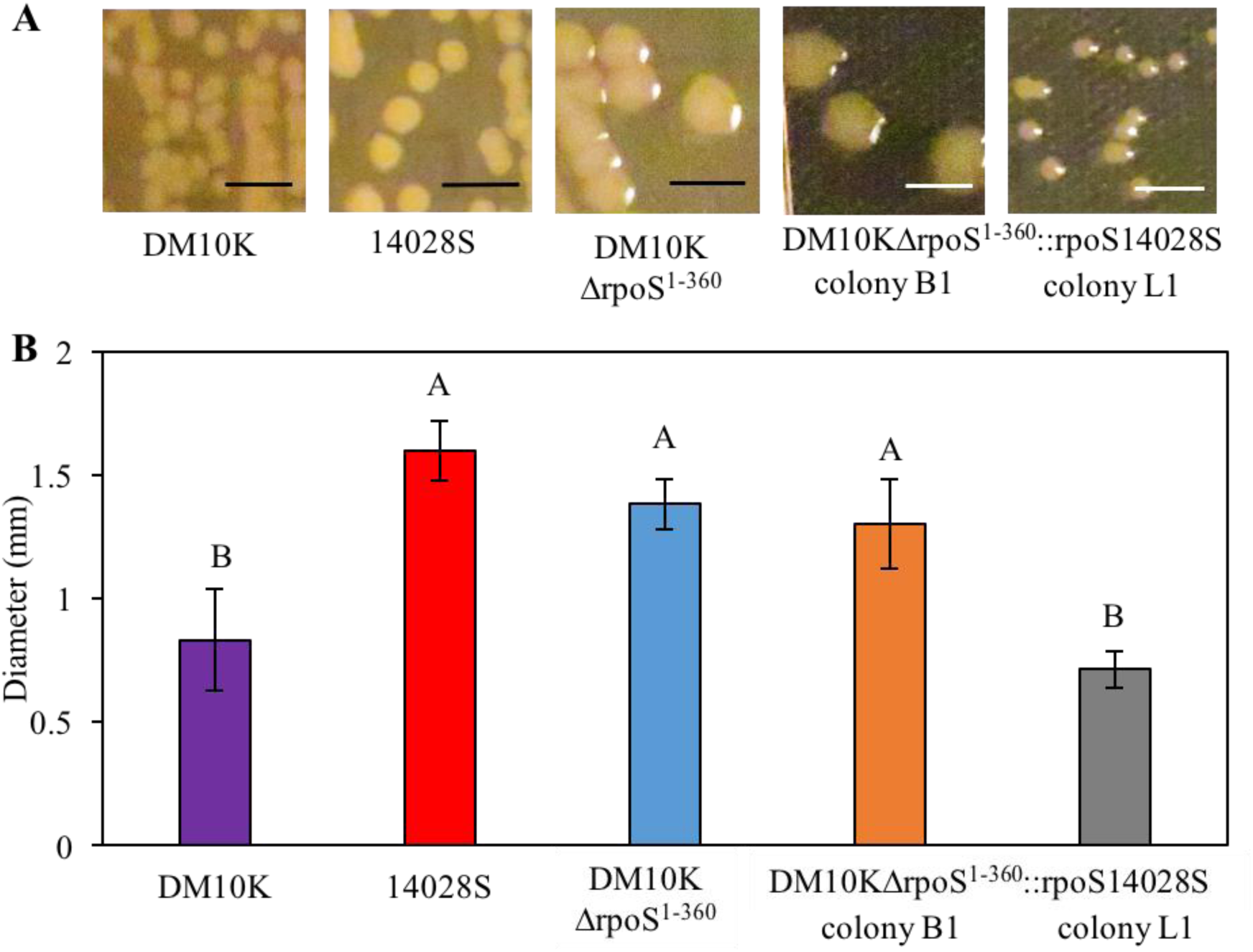
*S. enterica* DM10000 *rpoS* complement strains have variable colony size. **A**) Colony morphology and **B**) colony diameter of *S. enterica* strains 14028S (red), DM10000 (DM10K) (purple), DM10KΔrpoS^1-360^ (blue) and DM10KΔrpoS^1-360^::rpoS14028S colonies B1 (orange) and L1 (gray). Strains were streaked to isolation on LB plates from overnight cultures and incubated at 37°C overnight before imaged. Data are means ± SD (n = 5 colonies). Different letters indicate significant differences (1-way ANOVA for each inoculum at p < 0.05). Scale bar = 2 mm.

**Fig S7.**
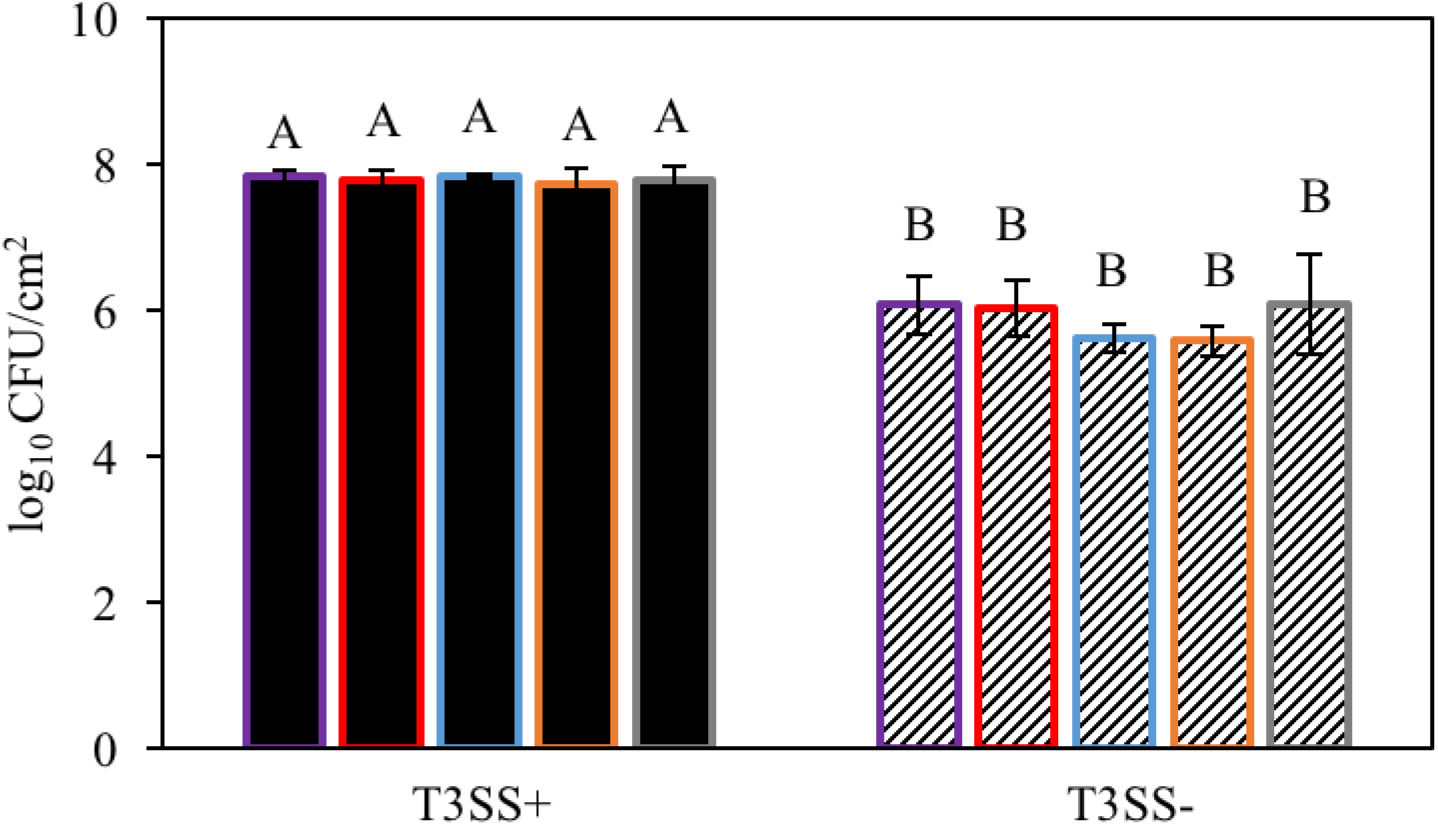
*P. syringae* causes disease in *N. benthamiana* during co-inoculation with all *S. enterica* strains. Bacterial populations of *P. syringae* pv. *tomato* DC3000Δ*hopQ1-1* with (T3SS+) and without (T3SS-) a functional Type III Secretion System after co-inoculation with *S. enterica* strains 14028S (red), DM10K (purple), DM10KΔrpoS^1-352^ (blue), and DM10KΔrpoS^1-352^::rpoS14028S colonies B1(orange) and L1 (gray). Inocula were syringe infiltrated into *N. benthamiana* at concentrations defined in Fig2. Data are means ± SD (n = 3 plants). Different letters indicate significant differences (2-way ANOVA at p < 0.05).

**Supplemental Table 1.**
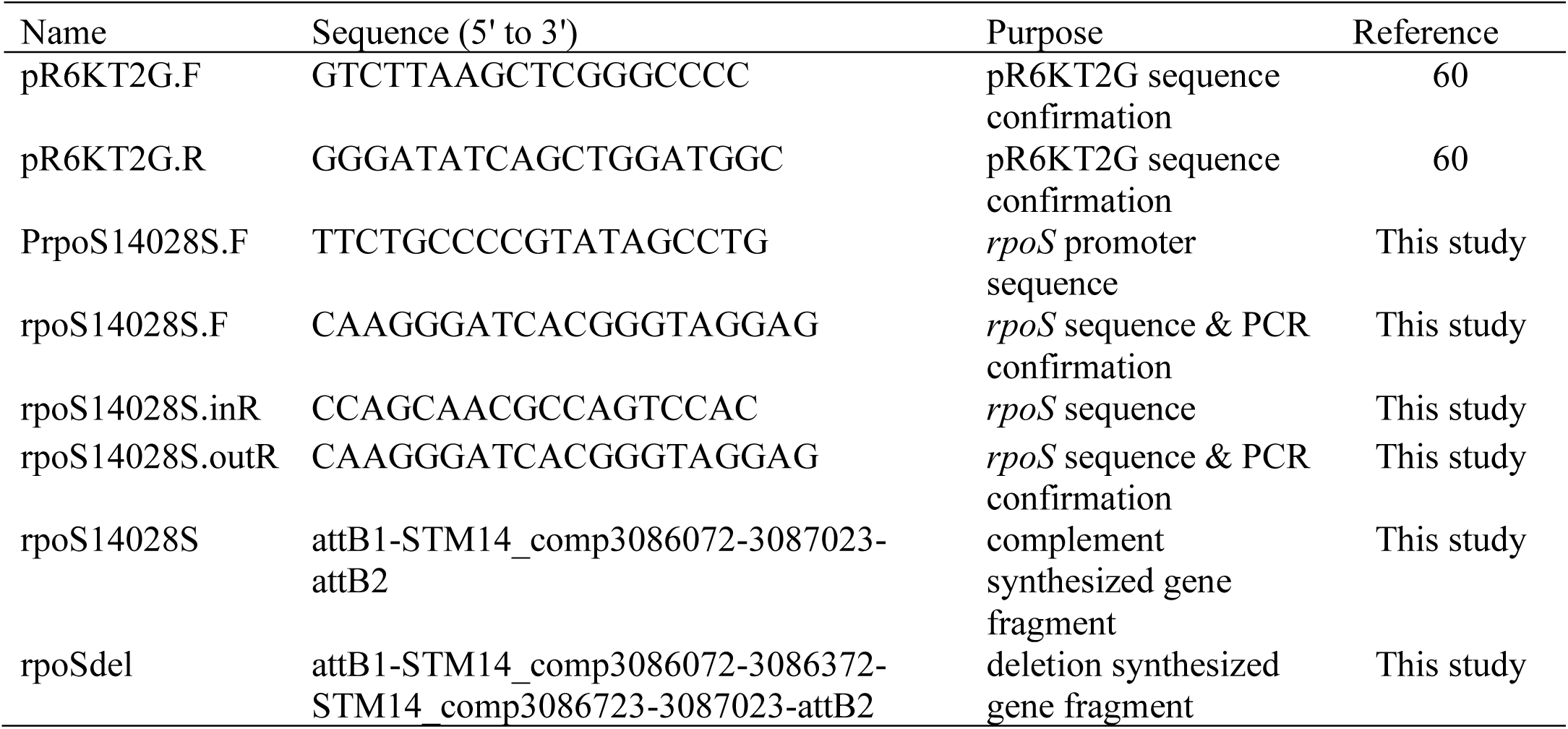
Primers used in this study.

**Supplemental Table 2.** GC-MS analysis of apoplast wash fluid (AWF) collected from *Nicotiana benthamiana*.

|  | Compound<sup>a</sup> | Relative<br>Concentration<sup>b</sup> | STDEV |
| --- | --- | --- | --- |
| 1 | Alanine (2TMS) | 149.04 | 36.93 |
| 2 | Glycine (2TMS) | 2.69 | 1.83 |
| 3 | Leucine (1TMS) | 7.95 | 1.08 |
| 4 | Proline (1TMS) | 345.54 | 256.22 |
| 5 | Valine (2TMS) | 27.99 | 3.09 |
| 6 | Serine (2TMS) | 43.53 | 12.80 |
| 7 | Leucine (2TMS) | 16.07 | 3.80 |
| 8 | Isoleucine (2TMS) | 12.85 | 1.81 |
| 9 | Threonine, DL- (2TMS) | 12.39 | 2.72 |
| 10 | Proline (2TMS) | 264.25 | 130.84 |
| 11 | Glycine (3TMS) | 6.03 | 3.56 |
| 12 | Serine (3TMS) | 43.46 | 16.90 |
| 13 | Threonine (3TMS) | 8.84 | 2.96 |
| 14 | Methionine (1TMS) | 0.09 | 0.07 |
| 15 | Aspartic acid (2TMS) | 32.55 | 21.74 |
| 16 | Asparagine [-H<sub>2</sub>O] (2TMS) | 1.33 | 0.47 |
| 17 | Aspartic acid (3TMS) | 151.93 | 39.88 |
| 18 | Pyroglutamic acid (2TMS) | 167.00 | 54.47 |
| 19 | Cysteine (3TMS) | 30.00 | 11.84 |
| 20 | Glutamic acid (3TMS) | 88.48 | 46.44 |
| 21 | Phenylalanine (2TMS) | 8.88 | 2.16 |
| 22 | Asparagine (3TMS) | 0.11 | 0.07 |
| 23 | Cysteinesulfinic acid (3TMS) | 0.00 | 0.00 |
| 24 | Glutamine, DL- (3TMS) | 3.23 | 1.34 |
| 25 | Arginine [-NH<sub>3</sub>] (3TMS) | 0.27 | 0.15 |
| 26 | Lysine (3TMS) | 1.64 | 2.38 |
| 27 | Tyrosine (2TMS) | 0.08 | 0.06 |
| 28 | Lysine (4TMS) | 1.44 | 0.34 |
| 29 | Histidine (3TMS) | 0.08 | 0.12 |
| 30 | Tyrosine (3TMS) | 6.79 | 1.32 |
| 31 | Histidine (4TMS) | 0.00 | 0.00 |
| 32 | Tryptophan, L- (1TMS) | 0.01 | 0.01 |
| 33 | Tryptophan (2TMS) | 0.02 | 0.00 |
| 34 | Tryptophan (3TMS) | 1.12 | 0.41 |
| 35 | Tryptophan (3TMS) | 1.11 | 0.42 |
| 36 | Cystine (4TMS) | 0.15 | 0.32 |
| 37 | Lactic acid, DL- (2TMS) | 0.10 | 0.14 |
| 38 | Glycolic acid (2TMS) | 0.14 | 0.11 |
| 39 | Oxalic acid (2TMS) | 0.10 | 0.04 |
| 40 | Malonic acid (2TMS) | 0.02 | 0.02 |
| 41 | Phosphoric acid (3TMS) | 3.12 | 1.58 |
| 42 | Nicotinic acid (1TMS) | 0.56 | 0.37 |
| 43 | Maleic acid (2TMS) | 31.46 | 9.88 |
| 44 | Succinic acid (2TMS) | 3.15 | 0.67 |
| 45 | Fumaric acid (2TMS) | 69.25 | 15.12 |
| 46 | Malic acid (3TMS) | 738.18 | 99.25 |
| 47 | Putrescine (3TMS) | 100.54 | 18.88 |
| 48 | Butanoic acid, 4-amino- (3TMS) | 459.55 | 82.01 |
| 49 | Glutaric acid, 2-oxo-1meox2tms | 0.39 | 0.60 |
| 50 | Tartaric acid (4TMS) | 0.00 | 0.00 |
| 51 | Putrescine (4TMS) | 12.58 | 11.66 |
| 52 | Shikimic acid (4TMS) | 0.03 | 0.01 |
| 53 | Citric acid (4TMS) | 24.02 | 6.44 |
| 54 | Quinic acid (5TMS) | 319.70 | 158.96 |
| 55 | Cinnamic acid,4-hydroxy,trans2tms | 0.00 | 0.00 |
| 56 | Glucose-6-p (1MEOX) (6TMS) | 0.00 | 0.01 |
| 57 | Glucose-6-P(1MEOX) (6TMS) | 0.00 | 0.01 |
| 59 | Rhamnose (1MEOX) (4TMS) MP | 61.71 | 0.90 |
| 60 | Fructose (1MEOX) (5TMS) MP | 130.89 | 82.83 |
| 61 | Fructose (1MEOX) (5TMS) MP | 158.50 | 25.30 |
| 62 | Glucose (1MEOX) (5TMS) | 181.08 | 101.02 |
| 63 | Glucose (1MEOX) (5TMS) MP | 210.75 | 54.83 |
| 64 | Galactose (1MEOX) (5TMS) | 210.75 | 54.83 |
| 65 | Galactose (1MEOX) (5TMS) MP | 42.03 | 13.42 |
| 66 | Glucose (1MEOX) (5TMS) | 210.75 | 54.83 |
| 67 | Mannitol (6TMS) | 55.37 | 32.33 |
| 68 | Inositol, myo- (6TMS) | 549.75 | 172.82 |
| 69 | Sucrose (8TMS) | 1272.19 | 96.53 |
| 70 | Trehalose, alpha,alpha'-, D8TMS | 0.00 | 0.00 |

<sup>a</sup> Compounds identified through GC-MS analysis of apoplast wash fluid collected from *N.*
*benthamiana* leaves. Derivatization was performed with Methyl-N-
(trimethylsilyl)trifluoroacetamide (MSTFA) and trimethylsilyl (TMS) derivatives are specified
for each compound.
<sup>b</sup> Area values of AWF compounds from six biological replicates were averaged and normalized
against an internal standard (ribitol) \* 100. Values are rounded to the nearest tenth.

